# Evolutionary jumps in bacterial GC content

**DOI:** 10.1101/2021.02.16.431469

**Authors:** Saurabh Mahajan, Deepa Agashe

## Abstract

Genomic GC content is a fundamental molecular trait linked with many key genomic features such as codon and amino acid use. Across bacteria, GC content is surprisingly diverse and has been studied for many decades; yet its evolution remains incompletely understood. Since it is difficult to observe GC content evolve on laboratory time-scales, phylogenetic comparative approaches are instrumental; but this dimension is rarely studied systematically in the case of bacterial GC content. We applied phylogenetic comparative models to analyse GC content evolution in multiple bacterial groups across two major bacterial phyla. We find that GC content diversifies via a combination of gradual evolution and evolutionary “jumps”. Surprisingly, unlike prior reports that solely focused on reductions in GC, we found a comparable number of jumps with both increased and decreased GC content. Overall, many of the identified jumps occur in lineages beyond the well-studied peculiar examples of endosymbiotic and AT-rich marine bacteria, and do not support the predicted role of oxygen dependence. Our analysis of rapid and large shifts in GC content thus identifies new clades and novel contexts to further understand the ecological and evolutionary drivers of this important genomic trait.

## Introduction

GC content refers to the fraction or percentage of GC base pairs in a genome. The GC content of bacterial genomes varies from as low as ~13% for *Zinderia insecticola* (McCutcheon & Moran 2012) to as high as ~75% for *Aneromyxobacter dehalogenans* (Thomas et al. 2008). Moreover, across bacteria the GC content of 4-fold degenerate codon sites varies from 5% to 95%, i.e. almost no GC base pairs to only GC base pairs (Hershberg & Petrov 2010; Muto & Osawa 1987). Such differences in GC content profoundly affect critical components of the expression of genomic information, including the usage of different synonymous codons (Knight et al. 2001), tRNA pools and tRNA modifying enzymes (Diwan & Agashe 2018), and amino acids (Knight et al. 2001; Lightfield et al. 2011). Given its fundamental importance for the maintenance and transfer of genetic information, the diversity of GC content and its evolutionary determinants have been investigated for many decades (Sueoka 1961). In general, the GC content of a sequence must be determined by a combination of biases in the mutational process, biases in the fixation process (selection or recombination), and drift. Although these forces may act differently on different sequences within a genome, the GC content of different regions such as intergenic regions, RNA coding genes, and protein coding genes and different codon positions within them are correlated to each other (Muto & Osawa 1987; Brocchieri 2014; Raghavan et al. 2012; Zhu et al. 2010). Thus, the GC content of a genome can be considered a single trait evolving under a set of common evolutionary pressures.

It is now well accepted that mutations in most bacteria (and also archaea and eukaryotes) are biased towards AT (Hershberg & Petrov 2010; Hildebrand et al. 2010) and the actual GC content of most bacteria is typically higher than expected based only on this mutation bias. Thus, on top of the underlying mutation bias, there is almost certainly also a fixation bias such that GC → AT mutations are preferentially removed or AT → GC mutations are favored. This fixation bias could be due to selection for higher GC content (Hershberg & Petrov 2010; Hildebrand et al. 2010), or due to a biased recombination process arising from GC-biased gene conversion (Lassalle et al. 2015). Although there are differences in the extent of the mutation bias such that mutations in AT rich bacteria are also more biased towards AT (Long et al. 2018), it is not clear if differences in the fixation bias contribute to GC content diversity. Additionally, a number of ecological factors have been proposed to be correlated with GC content (Agashe & Shankar 2014), e.g. host-association (Moran 2002), aerobiosis (Aslam et al. 2019; Naya et al. 2002), nitrogen fixation (McEwan et al. 1998), and temperature (Musto et al. 2004). However, many factors do not show strong correlations with GC content after accounting for the phylogeny or other confounding factors (Aslam et al. 2019; Marashi & Ghalanbor 2004; Vieira-Silva & Rocha 2008; Wang et al. 2006). Thus, the relationship between ecological factors and GC content diversity is also not clearly understood.

Since change in genome-wide GC content is a slow process, comparative analysis is by far the most informative approach to investigate the evolutionary determinants of GC content. Such studies of GC content diversity across bacteria have provided useful datasets and insights (Bobay & Ochman 2017; Hershberg & Petrov 2010; Hildebrand et al. 2010; Long et al. 2018). Across the range of GC content observed in bacteria, there are several trends characteristic of different bacterial groups. For instance, most Actinobacteria are GC-rich (average >60%), whereas Firmicutes are GC-poor (average ~40%) [for recent data, see (Lightfield et al. 2011; Reichenberger et al. 2015)]. Typically, closely related bacteria have similar GC content (Haywood‐Farmer & Otto 2003), although there are many well-studied exceptions. For instance, multiple lineages of insect endosymbionts and surface ocean dwelling bacteria have independently evolved exceptionally low GC content compared to their closest relatives (Giovannoni et al. 2014; Moran et al. 2008). Perhaps the most well-known example of the first kind are bacteria from the genus *Buchnera*, endosymbionts of aphids, whose average GC content is <30% compared to the ~50% GC content of related Enterobacteria (Chi-Yung & Baumann 1992; Moran 1996). Endosymbionts of many other insects also show similar trends of drastically reduced GC content (Moran et al. 2008). A well-known marine bacterium with exceptionally low GC content is *Pelagibacter*, an α-proteobacterium with a GC content of ~30% compared to ~50-60% of most other α-proteobacteria (Giovannoni et al. 2014). Similarly, other marine bacterial lineages are also highly AT-rich (Ghai et al. 2013; Giovannoni et al. 2014; Grzymski & Dussaq 2012; Luo et al. 2017). The extremely low GC contents of endosymbionts or AT-rich marine bacteria are clearly derived from a higher ancestral GC content (that is closer to their respective relatives) by drastic reductions in a relatively short time. The large changes in GC content of these lineages are explained by peculiar biological circumstances such as reduction in overall selection efficiency accompanying endosymbiosis (Moran et al. 2008; Wernegreen 2015) or intense selection associated with nutrient-poor surface ocean waters (Giovannoni et al. 2014). In contrast, closely related clades do not appear to have undergone such rapid changes in GC content. Thus, the evolution of GC content evolution appears to proceed differently in these special lineages *vis-a-vis* their relatives. However, it is not clear whether the distinct modes of evolution are a general feature of GC content diversification across bacteria.

Perhaps even more intriguingly, hitherto there are no reports of bacterial lineages with large increases in GC content. It is possible that such lineages exist, but have simply not been identified yet. On the other hand, large changes in GC may occur only in specific biological circumstances that cause reductions in GC content. These alternative scenarios have interesting implications for the diversity of GC content and its evolutionary drivers. If one were to find instances of large increases in GC, it would immediately raise many interesting questions. How frequently do they occur and in what biological circumstances? What are the evolutionary forces behind such changes, and are they similar to those causing GC reductions? Identification of such cases would also broaden the available datasets to better understand the evolution of GC content.

Questions about the generality of the different modes of GC evolution and its direction can be addressed using phylogenetic models of trait evolution (Felsenstein 1985; Pagel & Harvey 1989). These models are regularly used to study the evolution of morphological (Baker & Venditti 2019; Barkman et al. 2008; Landis & Schraiber 2017), behavioral (Hagey et al. 2017; Remeš et al. 2015) (Hagey et al. 2017; Remeš et al. 2015), and molecular traits (Liedtke et al. 2018; Stern & Crandall 2018) of animals or plants. Most simply, trait evolution on a phylogeny is modeled according to a Brownian process, i.e. as random changes accumulating at a constant rate without direction or constraint (Felsenstein 1985); or according to an Ornstein-Uhlenbeck process, i.e. as random changes occurring at a constant rate but with an attraction towards an optimal value (Hansen 1997). Modifications of these simple models capture more realistic evolutionary scenarios where the rate of trait evolution, the optimal value of a trait, or direction of evolution may differ across lineages (Beaulieu et al. 2012; Butler & King 2004; O’Meara et al. 2006). Another class of models based on the Lévy process capture qualitatively distinct evolutionary scenarios, where trait evolution is discontinuous due to occasional jumps in addition to accumulation of random changes at a constant rate (Duchen et al. 2017; Landis & Schraiber 2017). Comparison of the fit of different phylogenetic models and parameter variation across the phylogeny can provide interesting insights into the tempo and mode of trait evolution and their ecological and evolutionary correlates or mechanisms. Unfortunately, very few studies (Baidouri et al. 2017; Haywood‐Farmer & Otto 2003) have used such approaches to understand bacterial trait evolution. The evolution of the GC content of bacteria was previously analyzed using this approach (Haywood‐Farmer & Otto 2003), but before the advent of large datasets and sophisticated trait evolution models. This prior study found that GC content evolution is consistent with a Brownian model of evolution, implying gradual evolution at a constant rate. The discovery of bacterial lineages with rapid changes in GC content highlights the need for an expanded analysis with much larger and comprehensive datasets and new methods. Specifically, several phylogenetic models incorporating large jumps are now available and allow the inference of jumps in trait evolution (Duchen et al. 2017). These can be applied to large datasets of bacterial taxa to investigate GC content evolution.

We analysed the macroevolutionary patterns in bacterial GC content using such phylogenetic models. We found that GC content evolution is better explained by a combination of two modes of evolution: gradual diversification and relatively large “jumps”. Additionally, we identified specific branches that experience such jumps, and analyzed the ecological context in which they occur. We find that large changes in GC content are ubiquitous across bacteria, are not restricted to endosymbionts and marine lineages, and are not consistently related to changes in oxygen dependence. Interestingly, we found a large number of previously unrecognized instances of rapid increase in GC content. The macroevolutionary patterns found here raise further questions and provide interesting datasets to analyse the microevolutionary causes of GC content evolution in bacteria.

## Results

### A Lévy jumps model explains GC content evolution better than a Brownian model

As described in the methods section, we separately analysed GC evolution in 10 bacterial clades corresponding approximately to major orders from two large phyla (Table 1). For each order-level clade, we first evaluated and compared the likelihood of GC content distribution under a single-rate Brownian model and a single-optimum Ornstein-Uhlenbeck (OU) model. In all but two datasets, we found that the maximum likelihood estimate (MLE) of the constraint parameter (α) in the OU process was ~0 i.e. it just described a Brownian process without constraints. Moreover, in all cases the ML of the Brownian model was equal to the OU model (Table 1). Overall, GC content evolution was not consistent with constraint towards an optimal value. For this reason, we did not test the multi-optima OU models. Further, we found that in all cases, the Lévy jumps model (Duchen et al. 2017) explained the data significantly better than the single-rate Brownian model without jumps (Table 1). These results were consistent with our expectation based on the few known lineages with exceptional changes in GC content.

**Table 1:**
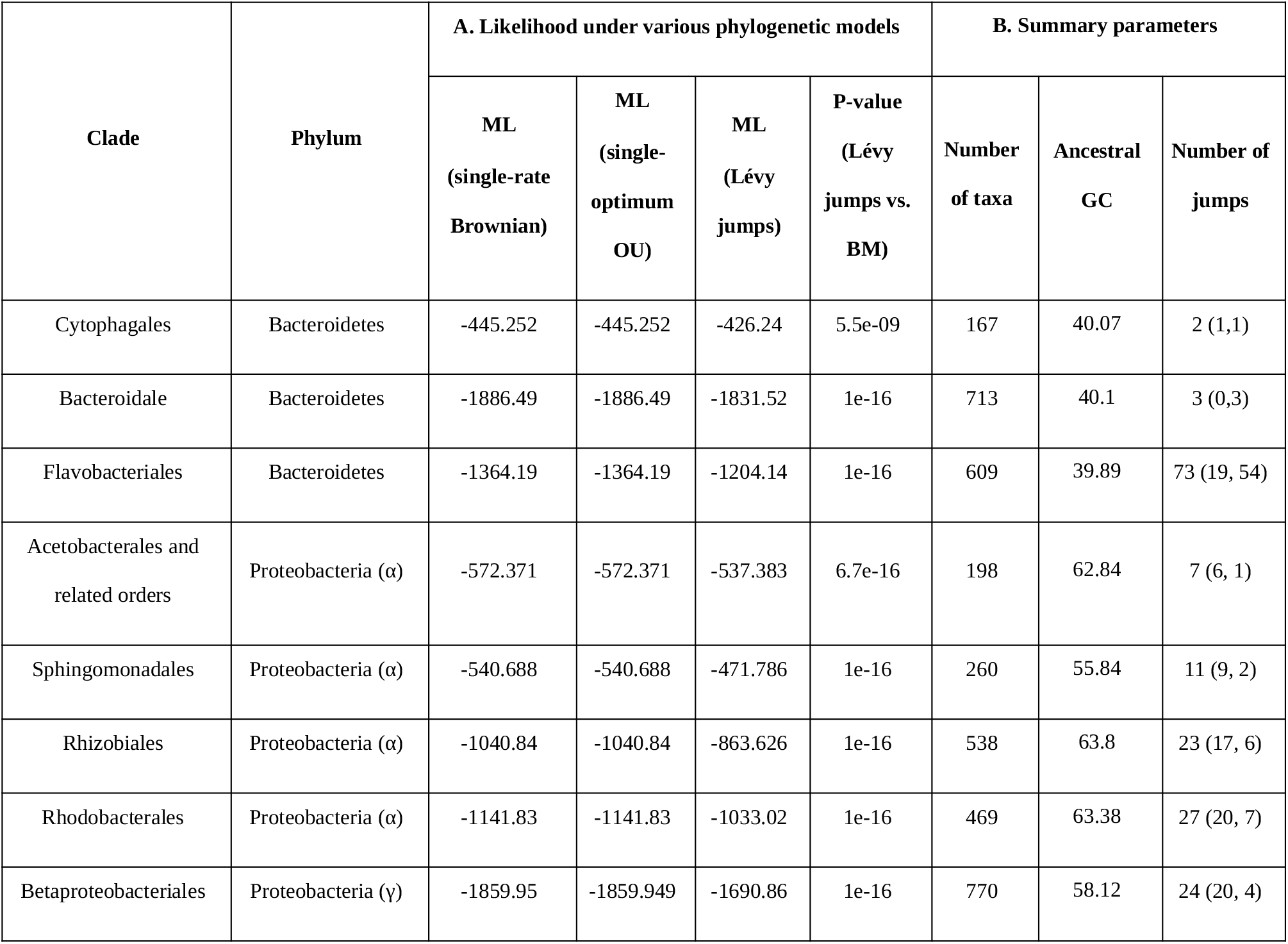
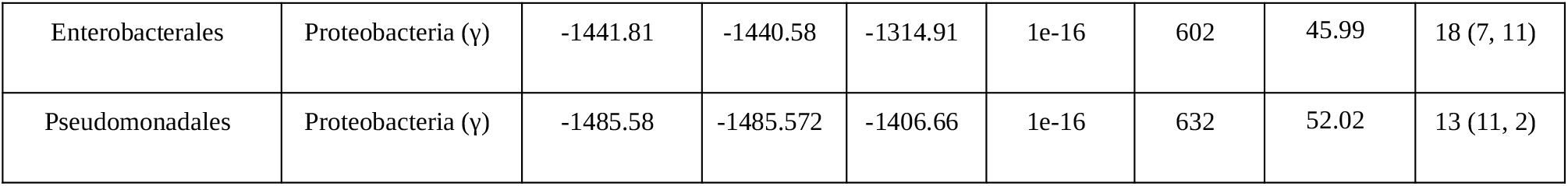
Summary statistics of phylogenetic models describing the evolution of GC content in various order-level bacterial clades. The table shows two sets of data for 10 order-level clades of bacteria: A. The maximum likelihood (ML) of GC content distributions under three phylogenetic models. The MLs of data under the Brownian and OU model are almost identical because the maximum likelihood estimate (MLE) of α, the constraint parameter in OU, was zero in almost all cases, making all best-fit OU models effectively equivalent to the Brownian models. The p-value of Lévy jumps model being better fit compared to the Brownian motion model was calculated from a likelihood ratio test (LRT). B. Some summary parameters. The number of jumps in the last column refers to the number of branches on which the posterior probability of detecting a jump was higher than the chosen threshold for each clade. As described in the methods, we treat each such branch as having experienced a single evolutionary jump in GC content. The numbers in parentheses refer to the number of downward and upward jumps, respectively.

| Clade | Phylum | A. Likelihood under various phylogenetic models |  |  |  | B. Summary parameters |  |  |
| --- | --- | --- | --- | --- | --- | --- | --- | --- |
|  |  | ML<br>(single-rate<br>Brownian) | ML<br>(single-optimum<br>OU) | ML<br>(Lévy<br>jumps) | P-value<br>(Lévy<br>jumps vs.<br>BM) | Number<br>of taxa | Ancestral<br>GC | Number of<br>jumps |
| Cytophagales | Bacteroidetes | -445.252 | -445.252 | -426.24 | 5.5e-09 | 167 | 40.07 | 2 (1,1) |
| Bacteroidale | Bacteroidetes | -1886.49 | -1886.49 | -1831.52 | 1e-16 | 713 | 40.1 | 3 (0,3) |
| Flavobacteriales | Bacteroidetes | -1364.19 | -1364.19 | -1204.14 | 1e-16 | 609 | 39.89 | 73 (19, 54) |
| Acetobacterales and<br>related orders | Proteobacteria ($\alpha$) | -572.371 | -572.371 | -537.383 | 6.7e-16 | 198 | 62.84 | 7 (6, 1) |
| Sphingomonadales | Proteobacteria ($\alpha$) | -540.688 | -540.688 | -471.786 | 1e-16 | 260 | 55.84 | 11 (9, 2) |
| Rhizobiales | Proteobacteria ($\alpha$) | -1040.84 | -1040.84 | -863.626 | 1e-16 | 538 | 63.8 | 23 (17, 6) |
| Rhodobacterales | Proteobacteria ($\alpha$) | -1141.83 | -1141.83 | -1033.02 | 1e-16 | 469 | 63.38 | 27 (20, 7) |
| Betaproteobacteriales | Proteobacteria ($\gamma$) | -1859.95 | -1859.949 | -1690.86 | 1e-16 | 770 | 58.12 | 24 (20, 4) |
| Enterobacterales | Proteobacteria ($\gamma$) | -1441.81 | -1440.58 | -1314.91 | 1e-16 | 602 | 45.99 | 18 (7, 11) |
| Pseudomonadales | Proteobacteria ($\gamma$) | -1485.58 | -1485.572 | -1406.66 | 1e-16 | 632 | 52.02 | 13 (11, 2) |

The estimated variance introduced by the Brownian component (σ_0_^2^), the jump rate (λ), and the total variance introduced by jumps relative to the Brownian component (λ·α) differed substantially across clades (Table S3, Supplementary File 2). The rate (or variance contributed per unit branch length) of the Brownian evolution component varied ~4 fold, with Flavobacteriales having the lowest and Bacteroidales the highest rate. The estimated jump rate varied >2 fold from ~2.5 jumps per unit branch length in Bacteroidales to >6 jumps in Flavobacteriales. The total variance introduced by the jumps per unit branch length was between ~2-fold lower (Bacteroidales) to >3-fold higher (Rhizobiales) than the Brownian component. Thus, the impact of the baseline (Brownian) rate as well as jumps in GC content evolution, both vary across bacterial orders. The reason for this variation in the frequency of jumps and the relative contributions of jumps to GC content diversity across clades is not very clear.

### Identification of branches experiencing GC content jumps

As pointed out earlier, using the procedure in *levolution*, one cannot predict the exact number or magnitude of jumps on each branch, but only estimate the posterior probabilities (pp) of the presence of >0 jumps (as defined in the model). Reliable inference of the phylogenetic location of jump(s) then requires one to choose an appropriate *pp* threshold. Since the jumps in the actual data are not known, we used simulated data to determine appropriate *pp* thresholds that led to optimal precision and recall in the inference of jump location. Briefly, we simulated GC content evolution according to the best-fit parameters of the Lévy jumps model, then inferred branch-specific posterior probabilities of jumps from the simulated GC contents of extant taxa, and determined the presence or absence of jumps on any branch according to a *pp* threshold. Varying this threshold and then comparing the inferred jump locations to the locations of simulated jumps allowed us to calculate precision and recall of jump detection under the different thresholds (see Methods for details).

In general, using high *pp* thresholds to infer jump locations leads to higher precision but poor recall, whereas using low *pp* thresholds to infer jump locations leads to lower precision but better recall in identifying branches with simulated jumps. However, the precision-recall relations of different order-level clades fell in two categories (Figure S1). For one set of clades, decreasing *pp* thresholds led to a small decrease in the precision as recall increased to ~30%, and a large decrease in precision thereafter. Sphingomonadales, Rhizobiales, Rhodobacterales, and Flavobacteriales are examples of this category. For a second set of clades, precision decreased rapidly and recall increased only slightly with decreasing *pp* thresholds. It is not clear why the precision-recall curves are different for these two categories, but may have to do with the specific tree topologies or taxon densities. Nevertheless, in the first case, we chose *pp* thresholds of 0.75 which lead to ~90% precision and ~30% recall in identification of simulated jumps. For the second case, we chose *pp* thresholds of 0.95 or 0.9 that also lead to ~90% precision but only ~2 to 20% recall. Although the recall appears very poor, larger jumps were detected more frequently as expected (Duchen et al. 2017). Simulated jumps with >5% GC content change had at least 40% recall, those with >10% GC content change had at least 60% recall, those with >15% GC content change had at least 80% recall, and finally those with >20% GC content change had almost 100% recall (Figure S2). Therefore, we expect that a majority of the branches experiencing large jumps in the actual data are identified correctly. These large jumps are also biologically more interesting and potentially insightful.

For inference of jumps in the actual GC content data, we identified branches with *pp* values greater than these thresholds (defined above) as those experiencing jumps. As examples, the inferred jumps mapped on a phylogeny of Rhizobiales and Acetobacterales are shown in Figure 1 and Figure 2 (for other clades, see Supplementary Figures S3-S10). Reassuringly, many instances of expected jumps in bacterial lineages with peculiar host-associated lifestyles were captured by this approach. For example, jumps were inferred at the stem branches for the Enterobacterial endosymbiont clades *Buchnera*+*Blochmannia* and *Baumannia*+others (Husník et al. 2011), at the base of a clade involving the Flavobacterial endosymbiont *Blattabacterium* (Bandi et al. 1995), and at the base of Betaproteobacterial (endo)symbionts *Kinetoplastibacterium* (Alves et al. 2013), *Polynucleobacter* (Heckmann & Schmidt 1987), and *Profftella* (Nakabachi et al. 2013). In Rhizobiales (Figure 1), a jump was inferred at the stem branch of the genus *Liberibacter*, which includes obligate host-dependent pathogens (Haapalainen 2014).

**Fig. 1.**
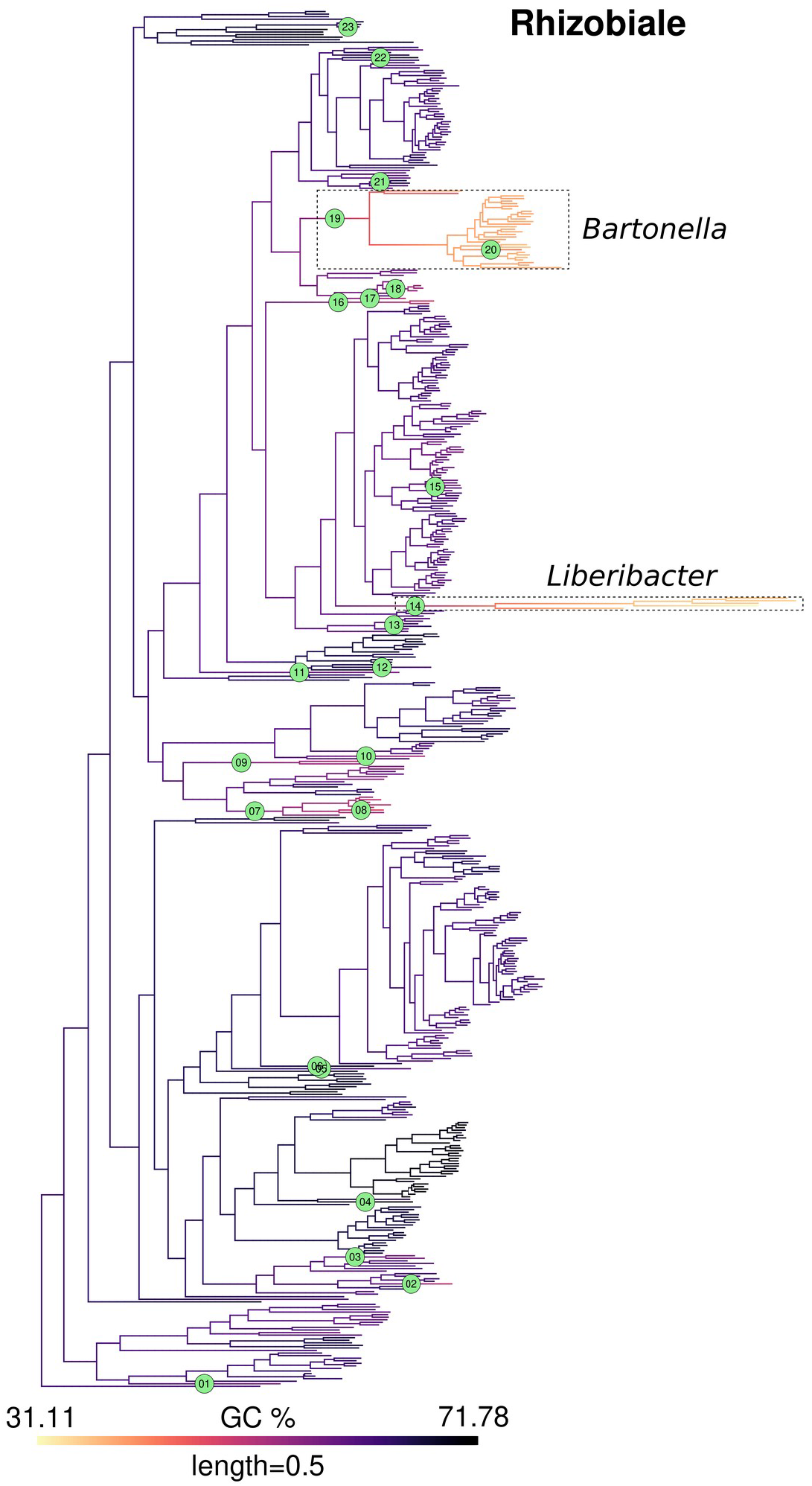
GC content map and location of inferred jumps in Rhizobiales. GC content was mapped onto a phylogeny of Rhizobiales using the contMap function from R package phytools. This mapping itself is only indicative of trends since it assumes a Brownian model of evolution. Branches with inferred jumps i.e. where the posterior probability of observing jump(s) is greater than the chosen threshold are indexed in green circles. Two interesting examples of jumps in Rhizobiales are highlighted in dashed boxes, which occur in the stem branches of *Liberibacter* (jump index 14), an obligate plant pathogen and *Bartonella* (jump index 19), an obligate animal pathogen, respectively. Within the genus *Bartonella*, the lineage leading to *B. australis* experienced an upward jump (index 20). Mapping for other clades is shown in Figure 2 and Figure S3 to S10.

**Fig. 2.**
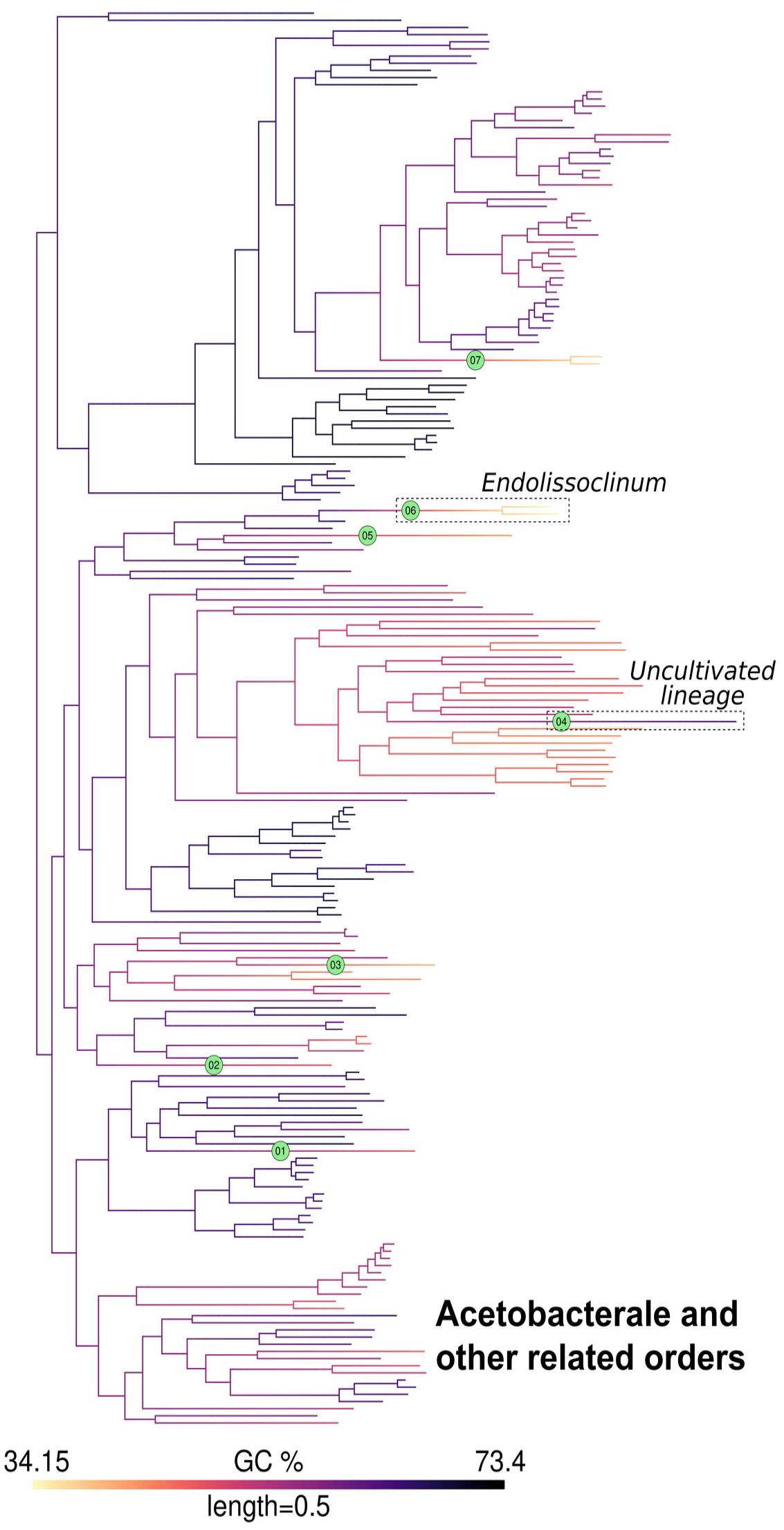
GC content map and location of inferred jumps in Acetobacterales and related orders. GC content was mapped onto a phylogeny of Acetobacterales and related orders as noted in Fig 2. Branches with inferred jumps i.e. where the posterior probability of observing jump(s) is greater than the chosen threshold are indexed in green circles. An example of a downward jump in an endosymbiont (*Endolissoclinum*, jump index 6) and an upward jump in an uncultivated bacterial lineage (jump index 4) are highlighted in dashed boxes. Mapping for other clades is shown in Figure 1 and Figure S3 to S10.

The total number of jumps inferred in this way ranged between 2 (for Cytophagales) to 73 (for Flavobacteriales) with a median of 15 (Table 1). Flavobacteriales appeared to be an exception since the next largest number of inferred jumps among other clades was 27 (for Rhodobacterales). However, the total number of detected jumps were not related to the ancestral GC content or the number of taxa in the clade (Table S3, Supplementary File 2). On the other hand, the fraction of upward jumps in a clade was related to its ancestral GC content. Clades with low ancestral GC content (<50%) experienced proportionally more upward jumps, whereas clades with high ancestral GC content (>50%) experienced more downward jumps (Figure 3A).

**Fig. 3.**
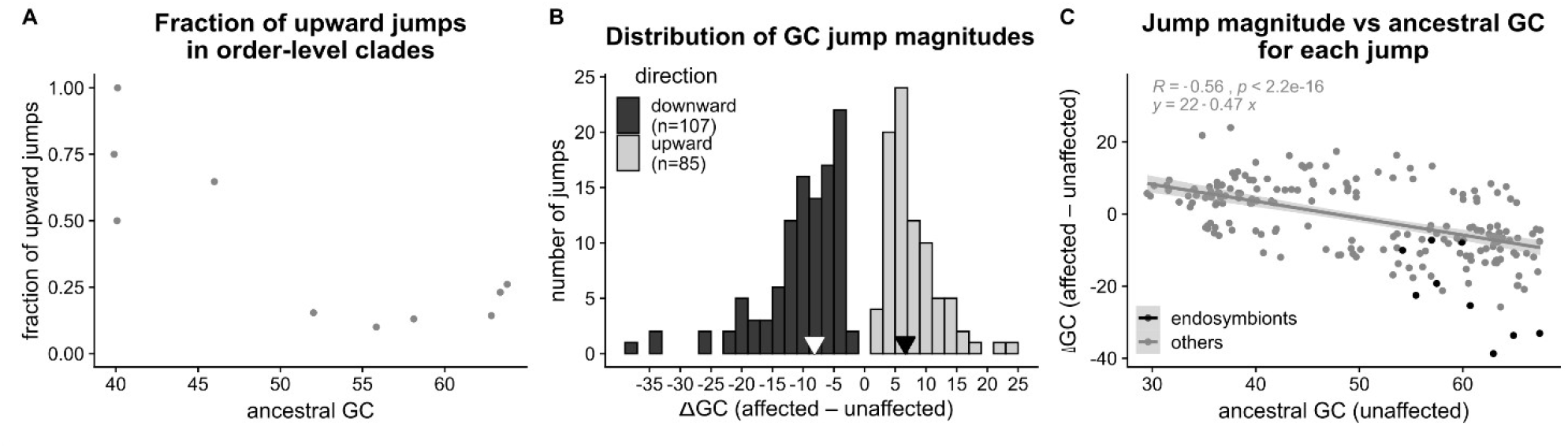
Direction and magnitude of GC jumps. We estimated the magnitude of each GC content jump as the difference in median GC content of descendant taxa of a branch affected by a jump (“affected”) and descendant taxa of a sister branch not affected by a jump (“unaffected”). Jumps involving increased GC content in affected taxa were designated as upward jumps, and those involving decreased GC content of affected taxa were designated as downward jumps. (A) The relation between the fraction of total jumps that were upwards and the ancestral GC content of each order-level clade. Ancestral GC content was estimated as a parameter of the Lévy jumps model using the procedure implemented in *levolution*. (B) Distribution of jump magnitudes. Black arrows denote the median magnitudes of upward and downward jumps. (C) Relation between jump magnitude and the estimated ancestral GC content, with the best-fit regression line (excluding non-endosymbiont clades). Ancestral GC content was estimated as the median GC content of unaffected taxa of the sister clade.

### Magnitude and direction of GC jumps

We estimated jump magnitudes as the difference between median GC contents of taxa affected by jumps, and taxa in the corresponding sister clade. Jumps occurred both in the upward (increasing GC%) and downward (decreased GC%) direction. Although downward jumps (n=107) were more frequent, we also found a comparable number of upward jumps (n=85) (Figure 3B). In terms of magnitude, downward jumps were bigger (ΔGC_median_ = −8.1%) than upward jumps (ΔGC_median_ = 6.6%). Even within the 10% largest jumps in each category, the average magnitude of the downward jumps was larger (ΔGC < −19.2%) than the upward jumps (ΔGC > 13.3%). Analyzed another way, among jumps with more than 15% change in GC, there were 18 downward jumps but only 5 upward jumps. Thus, while sudden increases in GC content are not rare, they tend to involve smaller changes in GC content compared to jumps that reduce GC%.

We further analyzed the direction and magnitude of jumps in relation to the estimated ancestral GC content (approximated as the GC content of sister clades that did not experience a jump in GC content). As expected, datasets with more extreme ancestral GC content (either lower or higher) were more likely to experience larger jumps in both directions (Figure 3C). The pattern was especially striking for endosymbionts with high-GC ancestors, which showed very large downward GC jumps.

Visually, it appears that jumps aree concentrated towards the tips i.e. towards more recent branches. However, this could also simply be a result of the number of branches being higher towards the tips in any phylogeny. Indeed, when we compared the distribution of the inferred jumps with jumps randomly placed on the phylogenies (with probability proportional to branch length), we observed that the distributions are not different (Figure S11).

### Ecological features associated with inferred GC jumps

Where possible, we extracted information from primary literature about the isolation source of taxa affected by the inferred jumps (“affected”) and closely related taxa unaffected by the jumps (“unaffected”) (Table S2, Supplementary File 2). Since we were interested in understanding the relationship between GC content jumps and *changes* in habitat or life history, we excluded clades where the isolation sources of both sets of taxa could not be reliably inferred. Of the 91 jumps where such data was available, 48 experienced decreased GC content and 43 experienced increased GC content (Figure 4).

**Fig. 4.**
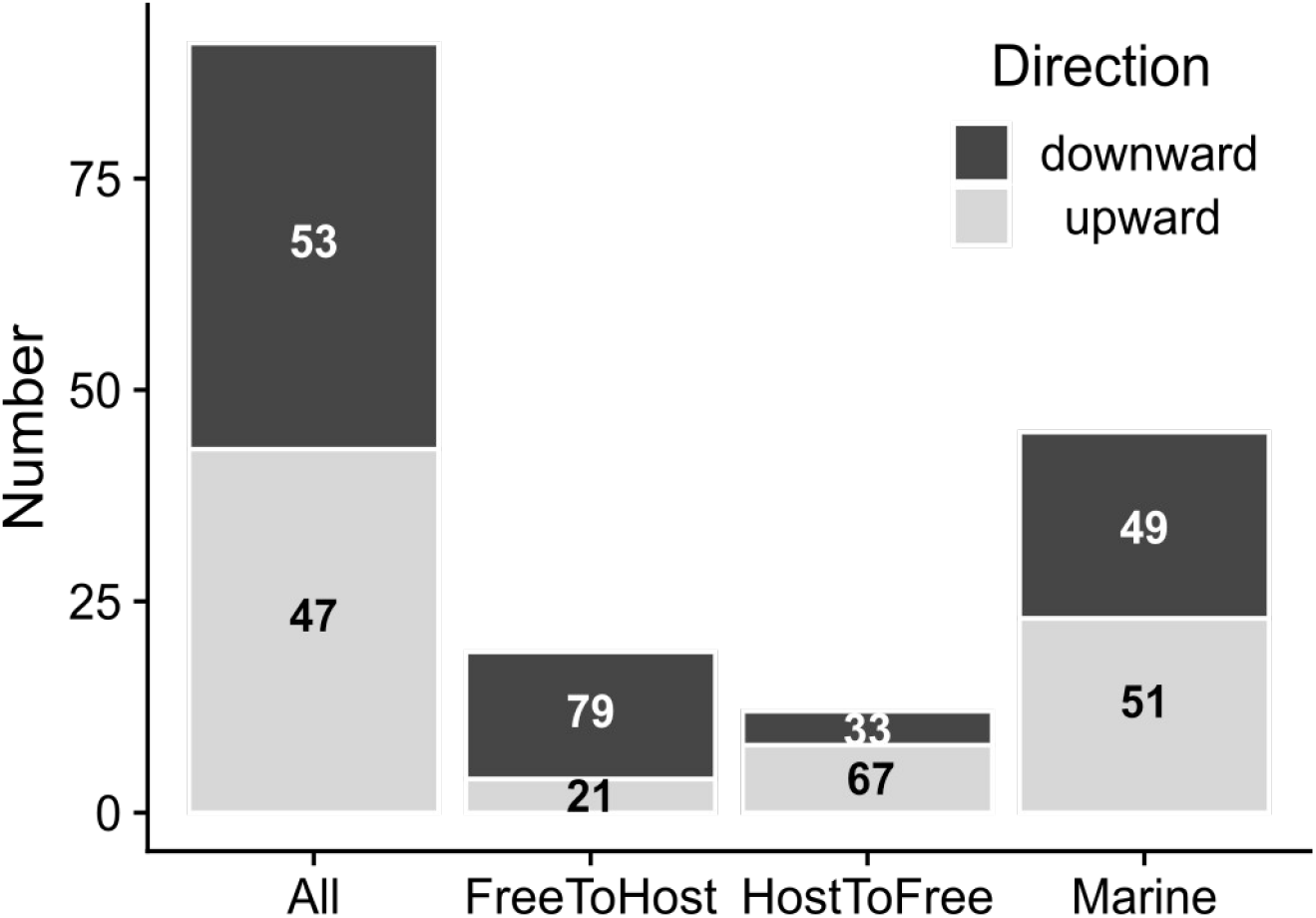
Proportions of upward and downward jumps across habitat and lifestyle categories. The number of upward and downward jumps are shown across 4 categories of datasets: (i) all datasets that could be analyzed for habitat or lifestyle related changes (n=91) (ii) a subset of datasets where the affected taxa (where a GC jump occurred) were host-associated, but related unaffected taxa were free-living (n=19) (iii) a subset of datasets where the affected taxa were not associated with hosts, but related unaffected taxa were host-associated (n=12), (iv) a subset of datasets where the taxa affected by the jump were isolated from marine habitats (n=45). Numbers in the bars denote the percentage of upward and downward jumps within each category.

In 7 of these jumps, affected taxa were obligatorily dependent on a host, and only in 4 of these they had switched to obligate host-dependence from a host independent lifestyle (Table S4, Supplementary File 2). Thus, only a minor fraction of jumps analyzed here are likely to be caused due to a strict dependence on a host and accompanying changes in evolutionary parameters. In the remaining 84 jumps, the affected taxa could be cultured independently on laboratory media. But in a further 19 of these, affected taxa were associated with hosts (i.e. were isolated from hosts or host-associated material) but unaffected taxa were not; implying a switch from free-living to host-association. Of these, 15 experienced a decrease in GC content, whereas 4 experienced an increase in GC content (Figure 4). Twelve other cases involved an opposite switch (from host-association to no host-association), associated with a GC jump. In this set, 8 affected taxa experienced increased GC content and 4 experienced decreased GC content (Figure 4). Thus, while about half the analyzed jumps involved a decrease in GC% (48/91 i.e. 53%), decreased GC content is more prevalent in affected taxa that appear to switch from no host-association to host-association (15/19 i.e. 79%) and less prevalent (4/12 i.e. 33%) in affected taxa that appear to switch in the opposite direction (p<0.05 for a Fisher’s exact test). Overall, while GC content jumps may sometimes arise due to changes in evolutionary parameters corresponding to such changes in lifestyle, a significant fraction of GC jumps (61/91 i.e. ~67%) did not involve association with (or separation from) hosts.

Independently, in more than half of the analyzed cases (45 of 84, excluding the 7 obligate host-dependent cases), affected taxa were isolated from marine habitats; suggesting that marine environments may impose distinct selection pressures that are especially likely to drive rapid GC shifts. However, in contrast to previous reports of GC reductions from AT rich marine bacteria, in half of the cases (23 of 45) affected taxa isolated from marine habitats showed increased GC content (Figure 4). Although it was not always possible to discern the exact niche of the involved taxa, at least 12 were isolated from coastal sediments, unlike the AT-rich marine bacteria from nitrogen-poor surface ocean waters. Other taxa affected by large GC jumps were isolated from fresh water, soil, decaying wood, bioreactors, and fermented products (Table S4, Supplementary File 2)

Since oxygen dependence has been previously suggested to be associated with higher GC content in bacteria, we also determined changes in oxygen dependence among the clades affected by jumps. Within datasets where such information was available, affected taxa showed no difference in oxygen dependence compared to related unaffected taxa in the majority of cases (44 of 56 comparisons); whereas affected taxa had increased oxygen dependence (predominantly, a change from facultatively aerobic to aerobic) in 7 cases and decreased oxygen dependence (a change from aerobic to facultatively aerobic or anaerobic) in 5 cases (Table S5, Supplementary File 2).

Interestingly, in 6 of 7 cases where affected taxa had increased oxygen dependence, they had also experienced increased GC content. Overall, in ~10% of analyzed cases, affected taxa were more dependent on oxygen and experienced increased GC content.

## Discussion

### Ubiquitous evolutionary jumps in GC content across bacteria

In the work presented here, we analyzed the evolution of GC content across large bacterial datasets (10 orders across 2 bacterial phyla) using phylogenetic models of trait evolution. We found that the diversification of bacterial GC content is more consistent with a mixture of Brownian evolution and ubiquitous evolutionary jumps, rather than pure Brownian evolution. As indicated by the previously reported examples of large evolutionary decreases in the GC content of endosymbionts and AT-rich marine bacteria, evolutionary jumps in bacterial GC content were not entirely unexpected. However, we find that the estimated variance in GC content contributed by such evolutionary jumps was more than the estimated variance contributed by the Brownian component in almost all bacterial clades that we analyzed. Thus, evolution by jumps appears to make a major, but thus far unrecognized, contribution to the diversification of bacterial GC content. Our results are also supported by another recent study that found pulsed evolution to be common in bacterial genome traits, including GC content (Gao & Wu 2021). Our study further describes the frequency, magnitude, and phylogenetic context of the observed jumps in GC content. Specifically, we emphasize two novel observations about the characteristics and ecological context of GC content evolution. One, we find a large number of evolutionary jumps that increase GC content, in contrast to previous studies that exclusively report evolutionary reductions in GC content of bacteria. Two, we find evolutionary jumps in GC content that occur in ecological contexts beyond endosymbiosis and habitation of surface oceans.

Interestingly, we found a comparable number of jumps that increase or decrease GC content. However, upward jumps were not associated with a clearly identifiable set of lifestyles or habitats (such as endosymbiosis). Moreover, we did not find any lineages with large increases comparable to the large reductions in GC content of some of the endosymbionts, and upward jumps were smaller in magnitude than downward jumps on average. These patterns may explain the absence of prior studies recognizing such jumps, though a fifth of the identified upward jumps were moderately large (>10% increase in GC content). Although we have not attempted a detailed analysis of the ecological context or evolutionary causes of upward jumps, the identification of lineages experiencing such jumps presents an opportunity to study them in the future. We found that upward jumps were more common in datasets with lower ancestral GC content, whereas they were less common in datasets with higher ancestral GC content. This is consistent with an evolutionary constraint on the range of observed GC content across bacteria (~25% to 75%), and indicates that the constraint may also apply to evolutionary jumps.

We must highlight that better fit by a model compared to other competing models does not say anything about the absolute ability of the model to explain the data. A better fitting model among three poor models will still be a poor one. Therefore, one has to rely on independent tests of whether model assumptions are satisfied and whether the model offers a good explanation of the data. Unfortunately, such exact “goodness of fit” tests are not available for most macroevolutionary models (Pennell et al. 2015). Thus, it is not clear if the Lévy jumps model used here offers an adequate explanation of GC content macroevolution. The modest recall of jump locations simulated according to the best-fit model parameters raises some doubts about the adequacy of the Lévy jumps model. However, the inference of exact locations of jumps is performed separately from the calculation of overall likelihood of data given the Lévy jumps model and estimation of summary parameters such as average jump rate and magnitude. Even when jump locations cannot be efficiently inferred, the average rate and magnitude of jumps can still be estimated accurately (Duchen et al. 2017). Moreover, we found that larger jumps in our simulations were recalled with greater frequency, reaching perfect recall for jumps with more than 20% change in GC content. Hence, we suggest that the largest jumps in the evolution of GC content are likely accurately reflected in our analysis.

Our study here considers GC content as a single trait and uses general trait evolution models to reconstruct its evolutionary history. This approach is justified by the correlations between GC content of different genome features such as the 1^st^, 2^nd^, and 3^rd^ codon positions, genes with different expression levels, genic and intergenic regions etc. and common evolutionary forces such as mutation biases acting on this trait. However, ancestral reconstruction of complete gene sequences based on branch heterogeneous models may offer an additional source of information for reconstructing the evolutionary history of GC content. This approach benefits from the large number of available sites in genome data, but is challenging due to the computational complexity of sequence evolution models and uncertainties in the estimates of the large number of parameters. In the future, this approach could be gainfully applied on smaller datasets, perhaps those selected on the basis of the present study.

Additionally, we must acknowledge that the inference of jump locations is subject to the specifics of and uncertainties in the underlying phylogenies. For example, the phylogenies used here were derived after de-replication of available genomes, where a few representative taxa among a closely related set were retained. This is true for the derivation of the original datasets (Parks et al. 2018) as well as our study (see Methods). Such pruning may cause spurious jumps to appear if the retained taxa happen to have different GC content from the closely related taxa not represented in the phylogeny. However, this is unlikely to be true because the representative taxa either had high average nucleotide identity (ANI>90%) or belonged to the same species as the ones that were removed. Another major source of spurious jumps may be the uncertainties in branch lengths. Specifically, underestimation of branch lengths may lead to trait changes being identified as more exceptional than they are in truth. However, such uncertainties in branch length should not affect our analysis severely since the underlying phylogenies are based on a large number of genes. As discussed earlier, these uncertainties are also less likely to affect the inference of larger jumps in traits. Although Bayesian methods that account for uncertainty in tree topology and branch lengths would be ideal (Huelsenbeck et al. 2000), such methods are not available for the jumps model used in our analysis.

### Useful datasets for studying evolutionary factors affecting GC content

Which evolutionary factors lead to GC content diversification is still an unresolved question. Mutational biases correlate with GC content across a diverse set of bacteria (Long et al. 2018); thus, changes in mutational biases must contribute to changes in bacterial GC content. However, what causes changes in mutational biases across bacteria is itself not well understood. Deletion of specific DNA replication and repair enzymes alters mutation bias in some bacteria (Dillon et al. 2017; Foster et al. 2018; Weissman et al. 2019) and the natural loss of some repair enzymes is the most likely reason for changes in the mutation bias of endosymbionts (Moran et al. 2008; Wernegreen 2015). However, whether such loss or gain contributes to changes in mutation bias and GC content in other lineages has not been investigated so far.

The role of changes in selection or GC biased gene conversion (gBGC) in diversification of GC content is also not clear (Lassalle et al. 2015; Bobay & Ochman 2017). Reduced efficiency of overall selection in endosymbiotic bacteria must contribute to reduced selection for GC content, but whether similar changes contribute to other instances of GC change is unclear. Moreover, there are few compelling explanations about what aspects of the biology of organisms could influence these microevolutionary forces leading to GC content jumps. Many environmental factors such as growth temperature, oxygen requirement, and nitrogen availability have been proposed to affect selection on GC content; but none offer convincing evidence after accounting for phylogenetic relatedness in the datasets (Agashe & Shankar 2014). Our analysis of GC jumps also failed to offer strong support for a major role of these environmental factors.

Based on the insights provided from prior studies of endosymbionts and AT-rich marine bacteria, we surmise that jumps in GC content are more likely to be driven by large changes in one or few different evolutionary factors. In contrast, gradual diversification of GC content (or underlying factors such as mutation bias) across longer time-scales may be driven by smaller changes in a number of factors, making it difficult to clearly identify causal relationships. A recent study also proposes that sudden jumps in mutational biases that alter the direction of bias should be generally selectively favoured, because such shifts in mutation spectra can allow populations to access under-sampled mutational space (Sane et al. 2020). If true, this hypothesis may explain GC jumps involving both increase and decreases in GC content, without invoking specific selection pressures favoring a change in either direction. The bacterial lineages experiencing GC jumps identified here can serve as interesting datasets to test this hypothesis, as well as the role of specific evolutionary factors such as habitat, metabolic requirements, and DNA repair enzymes that may drive GC content changes.

While we think that the jumps in genome GC content are a result of changes in ecological and evolutionary forces acting on GC content *per se*, it is possible that the observed changes in GC content of a focal clade could have resulted from horizontal gene transfer of a significant number of genes from a host with different GC content. However, we found that the changes in genome GC content during jumps are also reflected in similar changes in median GC content of genes coding for ribosomal proteins (Figure S12) that are unlikely to be horizontally transferred.

### Habitats and lifestyles of clades experiencing GC jumps

In this study, we attempted a preliminary analysis of the ecological context in which GC content jumps occur. Ideally, one would like to statistically test the association between GC jumps and habitat changes. This requires a complete characterization of all instances of habitat change in the entire dataset, which in turn requires habitat data for all the hundreds of taxa included in this study. Since it was not possible for us to collect this data, we decided to only characterize the clades that experienced GC jumps. But even an analysis of the full set of inferred jumps was precluded by limited data availability. Some lineages were represented only by metagenome-assembled genomes, where lifestyle related information could not be obtained. Other lineages were not represented in systematic collections of microbial phenotypes, and hence we could not use them to analyse the impact of ecological factors. Overall, we could analyze less than half the datasets (n=91 out of 201) for lifestyle or habitat related information, and even fewer (n=56) for oxygen dependence. We hope that in future, new ecological data on some of the interesting lineages with large GC jumps will allow more robust analyses.

Although extreme GC changes in endosymbiotic bacteria are well-studied examples, only a small fraction (~7%) of the GC content jumps in our analysis were attributed to endosymbionts. However, this number is an underestimate for the following reason. In some cases, endosymbiont lineages of independent origins get erroneously lumped together as single clades due to long branch attraction. For example, *Buchnera* and *Blochmannia*, two endosymbiont genera with reduced GC content have independent origins (Husník et al. 2011), but appear as a single clade in the phylogenies used here. Consequently, our jump inference method detects a single jump (Enterobacterales, jump 15) at the stem of this clade instead of two separate jumps. However, such undercounting should have a small effect on the number of jumps involving endosymbionts, because not all jumps with endosymbionts involve multiple endosymbiont lineages.

Beyond endosymbionts, in a further ~25% cases, we found that GC jumps occurred in taxa that had either evolved towards or away from a host association. Changes to GC content in such cases may be explained by changes in evolutionary parameters accompanying changes in lifestyle (e.g. effective population size), but this explanation needs further investigation. Nevertheless, a majority of GC jumps (67%) do not involve a change in lifestyle with respect to host-association. Similarly, in terms of oxygen dependence, affected taxa in a majority of analyzed jumps (80%) did not show a change compared to related unaffected taxa; but in about 10% jumps, affected taxa were more dependent on oxygen (aerobic instead of facultatively aerobic) and had experienced increased GC content. This is consistent with some previous studies that found increased GC content to be associated with increased oxygen-dependence (Aslam et al. 2019; Naya et al. 2002). However, previous studies do not identify specific instances of such associations. The datasets identified here can allow a more detailed investigation of this association and the potential mechanism underlying it.

Separately, about half the analyzed GC jumps occurred in marine lineages and it is possible that streamlining selection in this habitat could be contributing to some of these GC jumps. It was not clear if these lineages were indeed from nitrogen-poor surface waters (as previously reported for AT-rich marine bacteria (Giovannoni et al. 2005, 2014; Luo et al. 2017)); but at least 20% were isolated from sediments. Previous studies find that the AT-richness of some bacterial lineages in surface oceans is part of a set of characteristics (genome reduction, smaller intergenic regions, increased coding density, fewer regulatory genes) attributed to streamlining selection due to nutrient limitation (Giovannoni et al. 2005, 2014; Grzymski & Dussaq 2012). The specific lineages identified in this study make it possible to assess whether streamlining selection may be relevant to the observed GC changes.

Finally, we find many instances of jumps in GC content of lineages neither related to hosts or marine habitats. As an outstanding example, *Zymomonas* (a genus of free-living, fermenting bacteria) have experienced a large reduction in GC content (~45%) compared to the sister genus *Sphingomonas* (GC content ~55 to 65%). We also identified jumps with >10% reduction in GC content of other putatively free-living bacterial lineages such as *Robiginitomaculum*+*Hellea* (marine), *Aquaspirillum serpens* (aquatic), *Janthinobacterium sp. B9-8* (soil), *Hirschia* (marine); and jumps with >10% increases in GC content of the lineages *Siphonobacter aqueclarae* (aquatic), *Ferrimonas* (sediment), and *Flavobacterium* CP2B (marine) (Table S4, Supplementary File S2). We hope that a detailed analysis of the relevant evolutionary factors in such datasets identified here would lead to further insights into the mechanisms of GC content evolution in bacteria.

## Conclusion

We analysed the diversity of bacterial GC content through a phylogenetic lens and found evolutionary jumps as a predominant mode of diversification of bacterial GC content. We identify these jumps as particularly interesting to study the ecological and evolutionary factors driving GC content evolution. We further surmise that evolutionary jumps – particularly those involving larger changes in GC content – could be driven by changes in ecological or evolutionary factors. However, we did not find strong support for any of the putative ecological factors previously implicated in GC content evolution. Since it will be difficult to experimentally study a large number of bacterial lineages, we suggest that immediate follow-up studies could focus on signatures of selection, drift, and ecological factors that could be gleaned from the available genome sequence data.

## Methods

The methods used in this study are summarized in Figure 5.

**Fig. 5.**
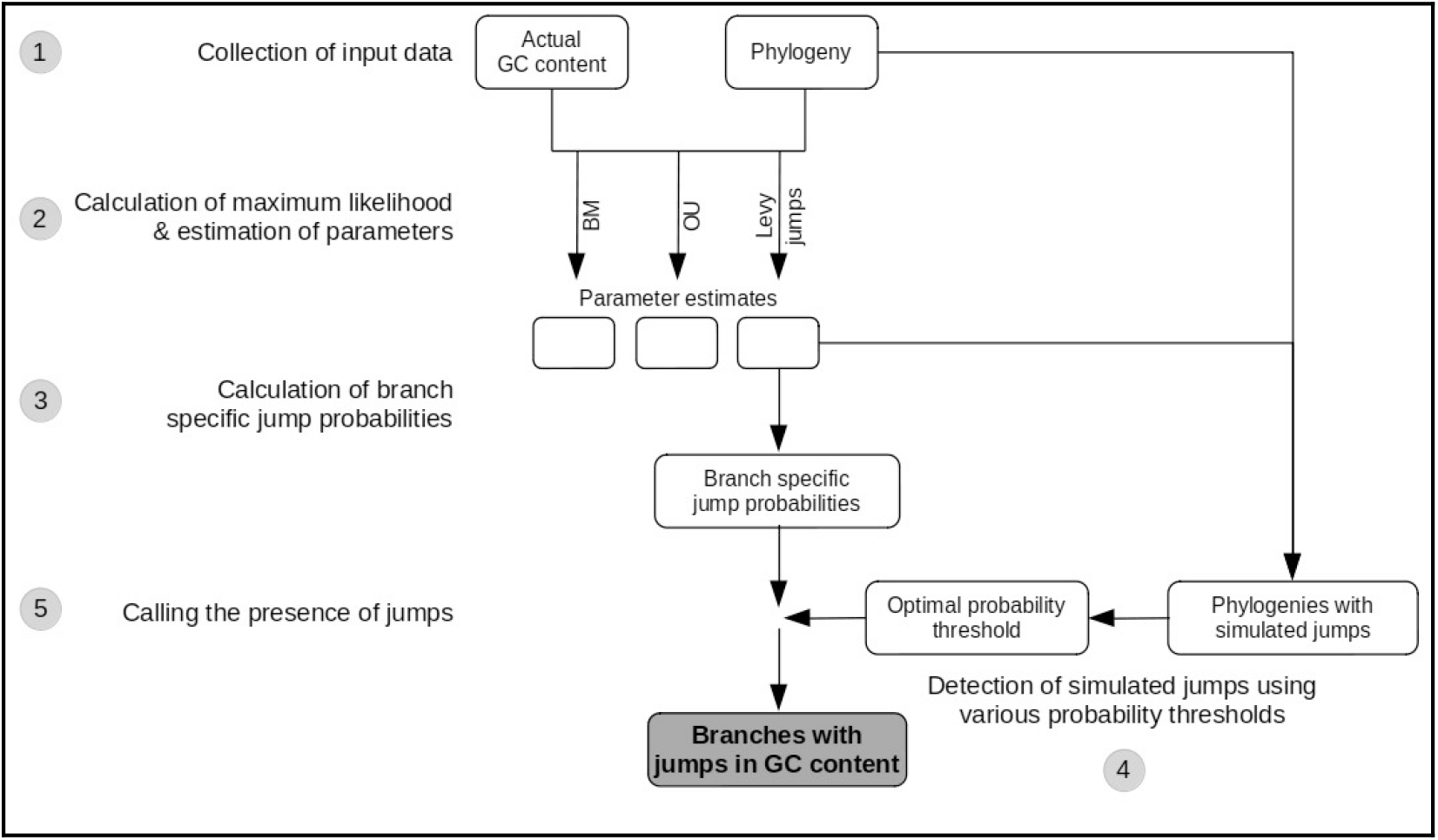
Summary of methods used in this study. Each major step in the analysis is numbered in the order in which it was performed. The analysis was performed independently for each of 10 order-level clades belonging to 2 bacterial phyla. Step 1: We derived the phylogenies of major bacterial clades and GC content of taxa from the Genome Taxonomy Database (GTDB). Step 2: We obtained the maximum likelihood (ML) and parameter estimates for different phylogenetic models using the phylogenies and the GC content distributions as the input. For the Brownian motion (BM) and Ornstein-Uhlenbeck (OU) models, these were obtained by exact analytical solutions implemented in the *geiger* package in R; while for the Lévy jumps model, these were obtained by Expectation-Maximization (EM) + Markov Chain Monte Carlo (MCMC) sampling method implemented in the *levolution* software. Step 3: For each branch in a phylogeny, we obtained the posterior probability of a jump in GC content using the phylogeny, GC contents, and the best-fit estimates of the parameters of the Lévy jumps model (obtained from step 2). These probabilities were obtained using an empirical Bayes approach implemented in the *levolution* software. Step 4: To calculate posterior probability thresholds to decide the presence or absence of jumps, we first simulated data with GC content jumps. The simulations were performed on the original phylogenies using the best-fit estimates of the parameters of the Lévy jumps model. We then attempted to detect the known jumps in simulated data using various posterior probability thresholds. We chose posterior probability thresholds that led to an optimal choice between precision and recall of the simulated jumps. Step 5: We deemed branches whose posterior probability of experiencing a jump (calculated from actual data in step 3) was greater than the optimal probability threshold (calculated from simulated data in step 4) as those having experienced a jump in GC content.

### Datasets

We started with a phylogeny of ~22,000 bacterial genomes inferred from universally conserved ribosomal proteins that was downloaded from the Genome Taxonomy Database (release 80) (Parks et al. 2018). The branch lengths of this phylogeny represent the number of substitutions per site. Due to uneven genome sequencing efforts, clades are unevenly sampled in this phylogeny; e.g. some bacterial species are represented by hundreds of genomes of different strains. These add redundant information about GC content diversity and may bias subsequent analyses. Therefore, we used a custom script to subsample and retain fewer representatives from densely sampled clades. Briefly, we scanned clades descending from every internal node, and retained only single taxa from clades younger than a specific threshold (0.01 substitutions per site). We chose this threshold in the following way. From the larger phylogeny, we first extracted a smaller clade containing Enterobacteria (which includes many densely sequenced species such as *Escherichia coli* and *Salmonella enterica*). We subsampled this clade with increasing threshold values retaining single taxa from clades younger than this threshold, and picked the threshold where the retained taxa consisted of only one or few strains from each named bacterial species.

To circumvent potential heterogeneity in the macroevolutionary process across distantly related branches and to reduce time required for downstream analyses, we extracted and analyzed subclades roughly at the level of taxonomic orders from the two largest bacterial phyla with genomic data: Bacteroidetes and Proteobacteria. Henceforth, the subclades are referred to as “order-level clades” and are specifically referred by the major taxonomic order contained in each of them. Across the two phyla, we analyzed 10 order level clades, each containing between ~200 to ~800 taxa.

We obtained genomic GC content data corresponding to all the analysed genomes from the Genome Taxonomy Database (release 80) (Parks et al. 2018).

### Comparing the fit of trait evolution models

We compared the likelihood of observed GC content distribution across the phylogeny, under three models of trait evolution: the single-rate Brownian (Felsenstein 1985), single-optimum Ornstein-Uhlenbeck (“OU”) (Hansen 1997), and the Brownian+stochastic jumps i.e. Lévy jumps (Duchen et al. 2017). The Brownian model describes a scenario where the trait value stochastically increases or decreases by a fixed amount per unit time, causing variance to increase at a constant rate (called the Brownian rate σ_0_^2^). The Ornstein-Uhlenbeck model includes an additional component that “pulls” the trait value towards an optimal value (parameter θ) with a speed that depends on a strength parameter (α) and the difference between the current and optimal value. A Lévy jumps model models a trait that evolves at a constant rate as in a Brownian model, but also experiences additional evolutionary changes as discrete and stochastic events (called “jumps”). In the model formulation used here (Duchen et al. 2017), these jumps are assumed to occur according to a Poisson process with frequency λ across the phylogeny. The average magnitude of jumps (i.e. changes in trait value) is modeled as a multiple (α) of the Brownian rate of evolution, although individual jump magnitudes are drawn from a normal distribution [Appendix I of (Duchen et al. 2017)].

We calculated the likelihood of data under the Brownian and OU models and the corresponding best-fit parameter estimates using the geiger package in R (Harmon et al. 2008). For the Lévy jumps model, we obtained the likelihood and the best-fit values of all parameters (except α) using an Expectation Maximization+Markov Chain Monte Carlo procedure implemented in the *levolution* software (Duchen et al. 2017). However, this procedure cannot directly estimate the maximum likelihood value of α, making it necessary to calculate the likelihood and estimate other model parameters independently for a range of α values, and then to choose an α that results in maximum likelihood. For this purpose, we evaluated α values in the set 0.1, 0.25, 0.5, 1, 2, and 4, i.e. letting the variance contributed by a jump vary between ten times less and four times more than the Brownian rate. From these separate calculations, we chose the α value and corresponding parameter estimates that resulted in the largest maximum likelihood (ML). In all the clades, ML values peaked in the range of α that were evaluated.

### Identifying evolutionary jumps in GC content

Given the estimated parameter values of a Lévy jumps model, one can also infer the phylogenetic location of jumps using the procedure implemented in the *levolution* software. This is accomplished by scanning multiple combinations of putative jump locations (branches) and then evaluating the posterior probabilities (*pp*) of one or more jumps occurring on every branch using an empirical Bayes approach (Duchen et al. 2017). There are two important issues that should be noted here. First, this procedure only allows the calculation of the posterior probability of a branch experiencing *one or more jumps* as defined in the theoretical model (Poisson events that introduce a specified amount of change in the trait value). This implies that one cannot know the exact number of theoretical jumps that are likely to have occurred on that branch. However, multiple theoretical jumps on a branch can be considered empirically equivalent to a single, but larger evolutionary change on the same branch. Therefore, subsequently in this study, we refer to the cumulative evolutionary change occurring on a single branch as a GC content jump. The second issue is that inferring jump locations is not a matter of yes or no, but of choosing an appropriate *pp* threshold above which we can reliably call a branch as having experienced a jump in trait value.

Ideally, all branches that have experienced a jump in trait value must have *pp*≈1 and all others, *pp*≈0. Thus, a high *pp* threshold should capture most jump locations accurately. In reality, this is not the case. Especially when the magnitude of jumps is small compared to the Brownian component, many branches that have experienced a jump have *pp* values much lower than 1 (Duchen et al. 2017). Therefore, choosing a high *pp* threshold can lead to a low “recall” of actual jumps. On the other hand, lowering the *pp* threshold to capture all jumps selects for many branches that have not actually experienced a jump (and therefore rightly received lower posterior probabilities). This lowers the “precision” of jump inference. Altogether, as the *pp* threshold is varied, there is a negative relationship between precision and recall.

Hypothetically, if the real jumps in any evolutionary history were known, one could choose a *pp* threshold that optimized precision and recall of the inference procedure. Of course, we do not know the location of actual jumps in GC content. Therefore, we determined *pp* thresholds that optimized both precision and recall of jump inference in simulated data with exactly known jumps.

For each order level clade, we decided *pp* thresholds in the following manner.

1. We modified the *ex.jumpsimulator()* function in the *geiger* package (Harmon et al. 2008) to simulate 5 independent datasets. In each simulation, GC content evolved according to a Lévy jumps model (described in the previous section) i.e. continuously changing according to a Brownian process with additional changes (“jumps”) at branches selected stochastically according to a Poisson process. The parameters for the model: ancestral GC content, Brownian rate (σ_0_^2^), jump rate (λ), and the average relative jump magnitudes (α), were set to the best-fit estimates obtained from actual GC content data of the respective clades. The location of each simulated jump was recorded by the function used to simulate the datasets.
2. Using the phylogeny of the clade and the simulated GC content of only the tips as inputs, we followed the procedure in *levolution* (explained in the previous section) to estimate the branch-specific posterior probabilities of jumps for each simulated dataset.
3. Independently for each *pp* threshold in a range of putative *pp* thresholds between 0 and 1, we calculated precision and recall in the following way:

3.a We inferred jumps in branches with *pp* higher than the threshold under consideration.
3.b We divided the inferred jumps into two types: “true jump estimates”, when a jump was inferred on a branch with a simulated jump; and “false jump estimates” when a jump was inferred on a branch with no simulated jump.
3.c We pooled data across the 5 simulations and calculated the precision and recall as:

c.i precision = 100 *x* number of “true jump estimates”/(number of “true jump estimates” + “false jump estimates”)
c.ii recall = 100 *x* number of “true jump estimates”/(number of simulated jumps)

We chose a *pp* threshold that led to at least 90% precision while trying to achieve maximum recall (Figure S1). The chosen thresholds for different clades resulted in a precision between 91 to 97% and a recall between 2 to 37%. We deemed that all branches of an order-level clade with a posterior probability greater than the chosen threshold had experienced a jump in GC content. In every order-level clade, each such branch was assigned a unique serial number (referred to as “jump index”) for reference.

When precision and recall was calculated separately for each simulated dataset instead of the pooled dataset, the chosen thresholds resulted in at least 80% and upto 100% precision in all cases (data summary in Table S1, Supplementary File 2). The recall varied considerably across independent simulations: for clades with low overall recall, it varied from 0% to 12% across simulations; whereas for clades with modest recall, it varied from 25% to 42% across simulations.

### Analysis of inferred jumps

To analyze the directions and magnitudes of the inferred jumps in GC content, we resorted to an approximate calculation because the procedure in *levolution* cannot estimate the magnitude of jumps occurring on each branch. Therefore, we quantified the impact of each jump by comparing the median GC content of all descendant taxa of the branch affected by the jump with the median GC content of all descendant taxa of the corresponding sister branch. If any descendant branches were also affected by additional nested jumps, we removed the corresponding descendant taxa from the calculation.

To test how often GC content jumps were associated with endosymbiosis (or other forms of host association) or with marine habitats, we searched for primary literature describing the isolation of taxa in clades affected by GC content jumps and their sister clades. We specifically looked for evidence of whether the organism could be cultured independently of a host. If the primary source mentioned that the organism could be isolated and grown independent of the host, we tagged it as: “not host dependent”. If the organism was isolated from a host, then we tagged it as: “host associated”. In this analysis, we excluded clades represented only with metagenome-assembled genomes, those with large sister clades containing diverse species, and taxa whose phylogenetic placement was unreliable (Table S2, Supplementary File 2).

In addition, we also obtained data about the oxygen dependence (anaerobic, facultatively aerobic, aerobic, or obligately aerobic) of taxa from a recent compilation of bacterial phenotypes (Madin et al. 2020). We manually assigned oxygen dependence to entire clades (those experiencing GC jumps or sister clades) based on the oxygen dependence of the majority of taxa in each clade.

## Author Contributions

SM contributed to the design of methods and analysis, performed the analysis, interpreted results, and wrote the manuscript. DA conceived the study and contributed to the design of methods and analysis, interpretation of results, and writing.

## Acknowledgments

This work was supported by the Council of Scientific and Industrial Research India (fellowship “19/372624/SPMF 2013 Award” to SM), the DBT/Wellcome-Trust India Alliance (grant IA/I/17/1/503091 to DA), and the National Centre Biological Science (NCBS-TIFR), Bangalore. We thank Gaurav Diwan for contributing to the script for pruning phylogenies, and for useful discussions.

## Data Availability Statement

The data and code underlying this article will be available as Online Supplementary Files and a Github repository at [https://github.com/saurabh-mk/manuscript-bacGC]. Published sources of data used to derive data in this study are referenced at appropriate places in this article. The repository mentioned above contains the trees containing posterior probability of jumps on each branch that were generated as part of the study. Other data regarding habitats and lifestyles of bacterial taxa compiled as part of this work are available as Supplementary Tables in Supplementary File 2. The repository mentioned above includes code and instructions necessary to reproduce the analyses in this article.

