## Supplementary Figures for "Evolutionary jumps in bacterial GC content"

All supplementary tables can be found in Supplementary File 1.

**Table S1 Variation in precision and recall of detecting simulated jumps across 5 independent simulations.**

Trait evolution was simulated 5 times using maximum likelihood estimates of parameters obtained from actual data. The location of simulated jumps was known and was used to calculate precision and recall after the inference step.

**T****able S2 List of taxa affected by jumps that were excluded from some subsequent analysis.**

MAG refers to clades where all or majority genomes were derived from metagenomes. GTDB Taxonomy refers to the taxonomic assignments at the Genome Taxonomy Database, where MAG taxa can be distinguished based on their names containing “UBA”. GC heterogeneity, clade size, and number of nested jumps were assessed using a code, and clades were excluded based on criteria laid out in the Methods section.

**Table S3 Summary of habitat and lifestyles of non-MAG clades affected by jumps and corresponding sister clades****.**

Jump index refers to unique index for each jump within an order level clade. In the reference column, F1, F2 and so on refer to references for taxa belonging to focal clades (affected by jumps) and S1, S2 and so on refer to references for taxa corresponding sister clades. A score of 1 in columns 7-9 indicates affirmative. In the last (10th) column “0” refers to no switch, “1” refers to a switch from no detected host dependence to non-obligate host association, and “2” refers to a switch to obligate host-dependence.

**Table S4 Oxygen dependence in non-MAG clades affected by jumps and corresponding sister clades****.**

Datasets refer to orders and jump indices. Subsequent columns indicate GC of the focal clade and sister clade, difference in GC i.e. magnitude of the GC jumps, oxygen dependence of taxa in focal clade and sister clade, and the inferred change in oxygen dependence.

**Figure S1. The precision-recall relations for jump inference on simulated data.**

Precision and recall of the jump inference procedure was evaluated on 5 simulated datasets for each clade. The orange point and associated label denote the posterior probability threshold chosen to infer jumps in actual data. Alpha1 refers to clade involving Acetobacterales and other related orders of α-proteobacteria.

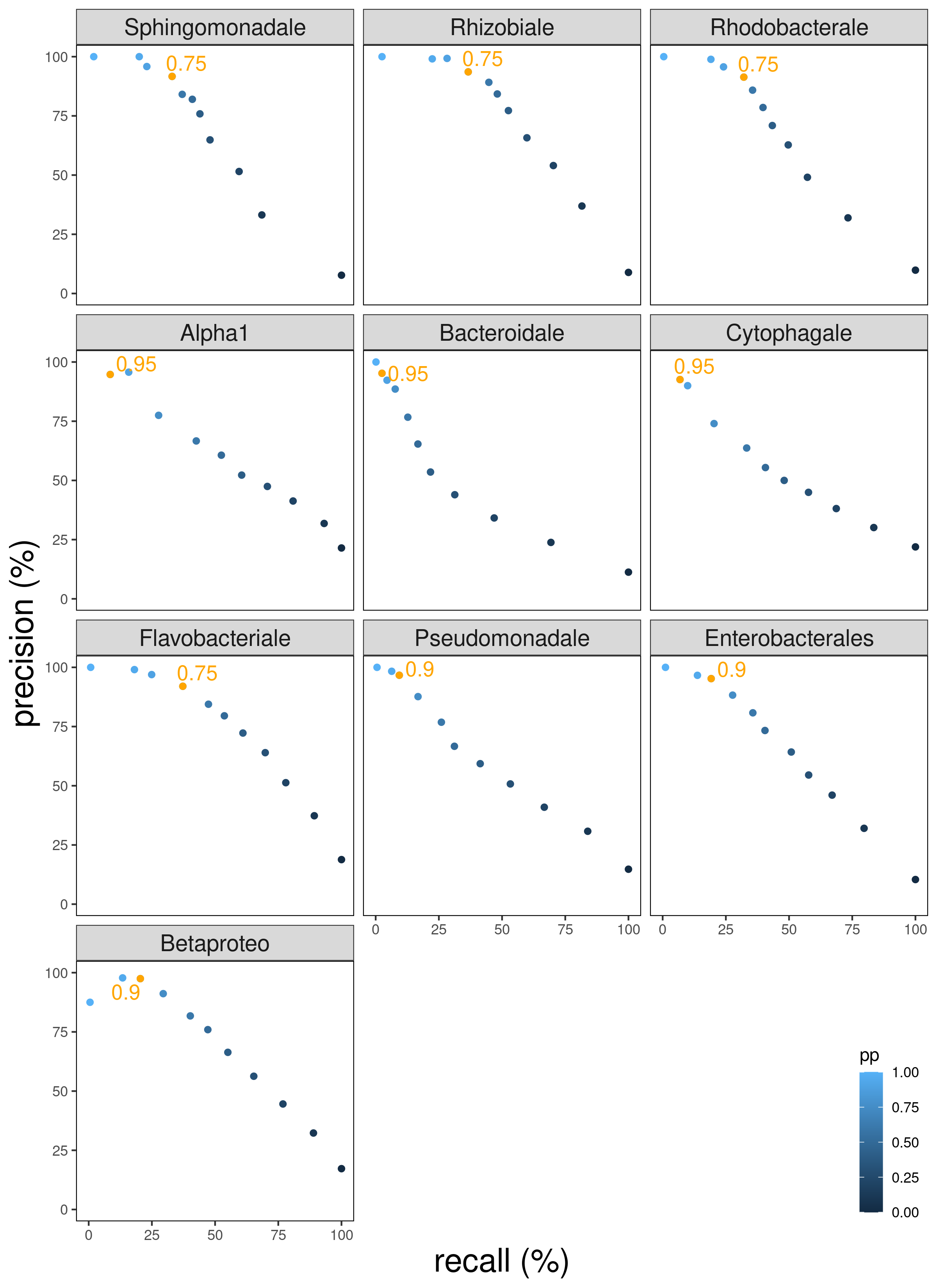

**Figure S2. Recall as a function of jump magnitude in simulated jumps.**

Simulated jumps across all clades and 5 replicate simulations each were pooled (total n~6000) and their magnitudes were obtained as the difference between GC contents of the descendant and ancestral nodes. Jumps were divided according to their magnitude in intervals of 5% change in GC (range -25% to 25%). Fraction of the simulated jumps that were successfully inferred (recalled) according to the procedure in *levolution* were then calculated for each set.

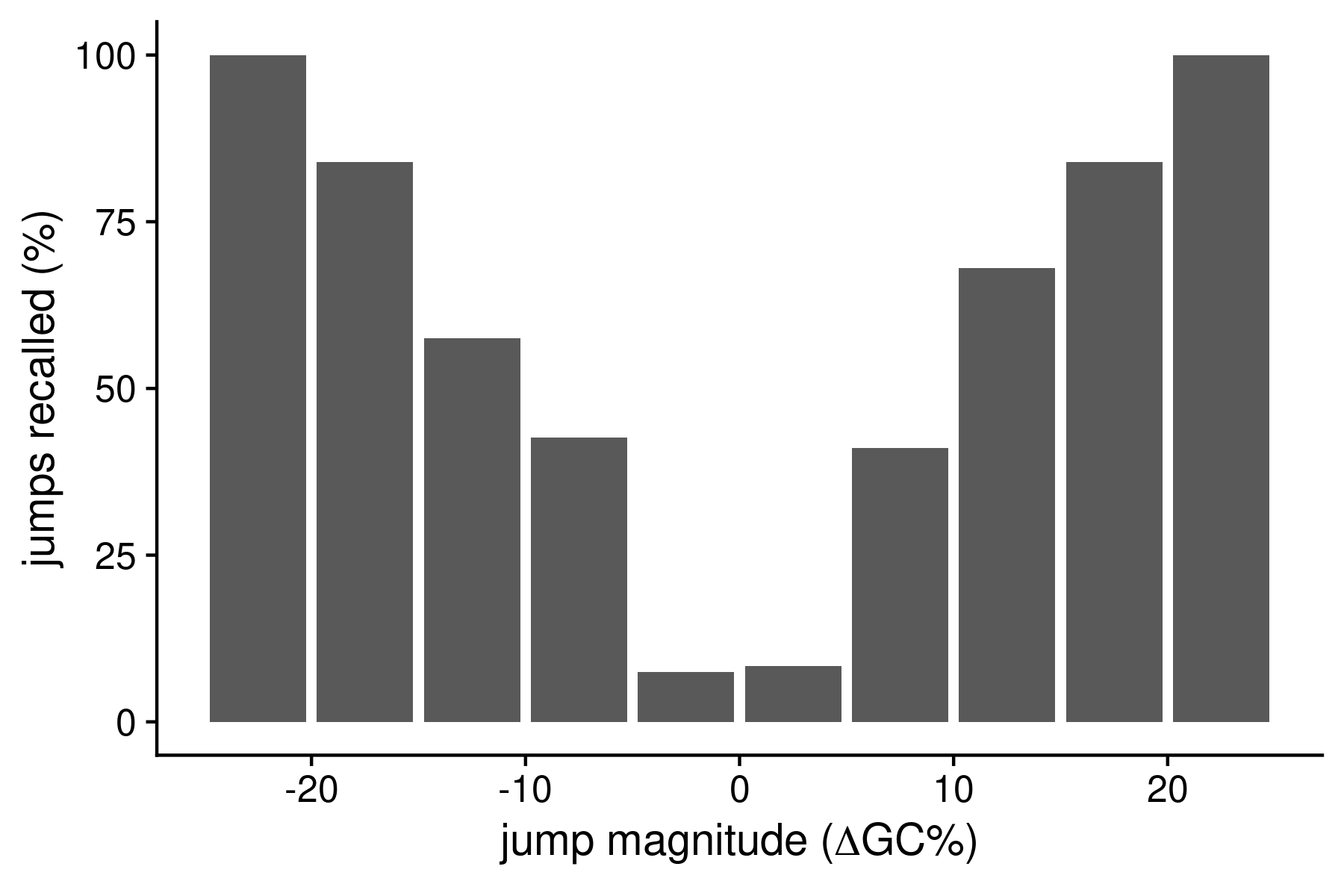

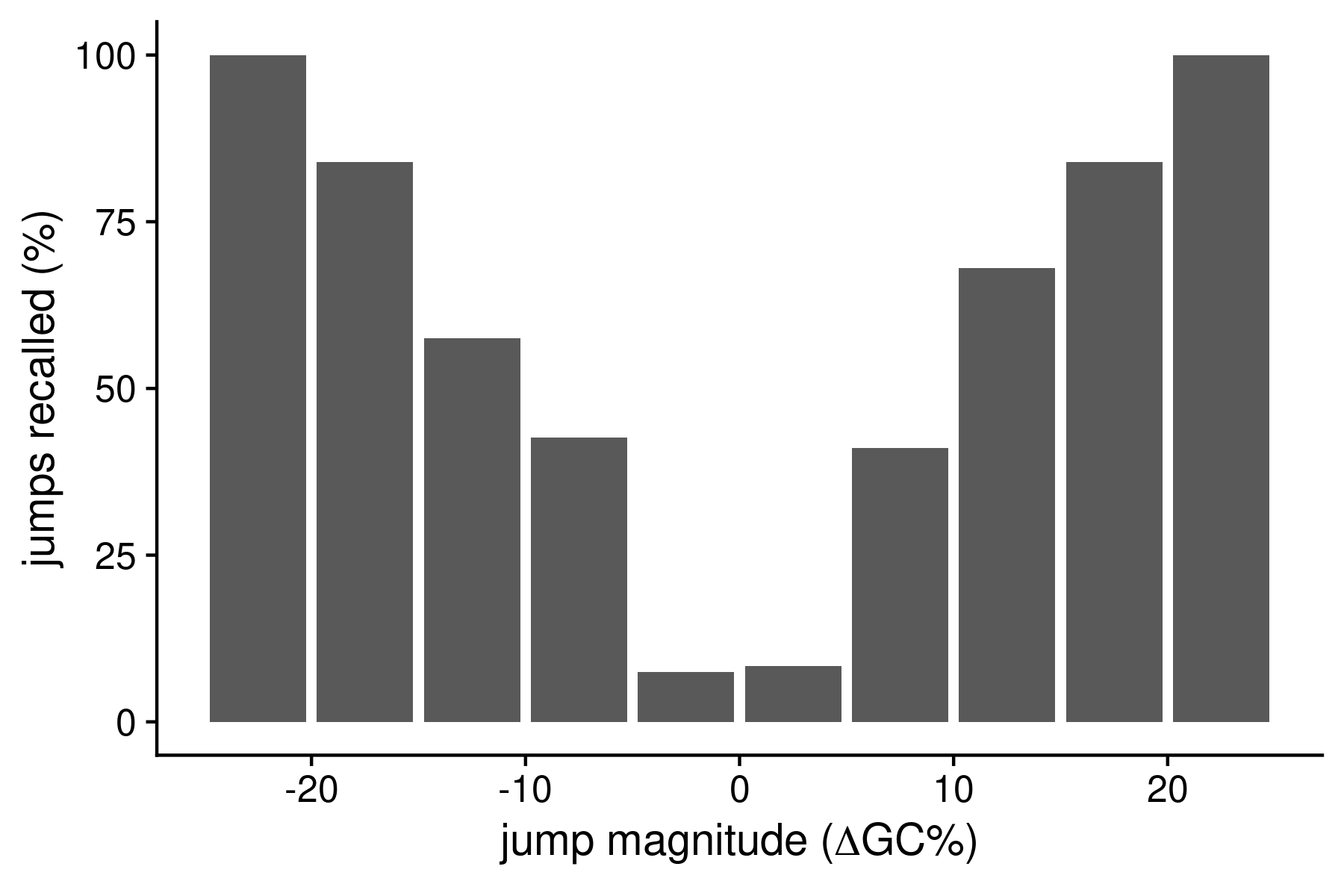

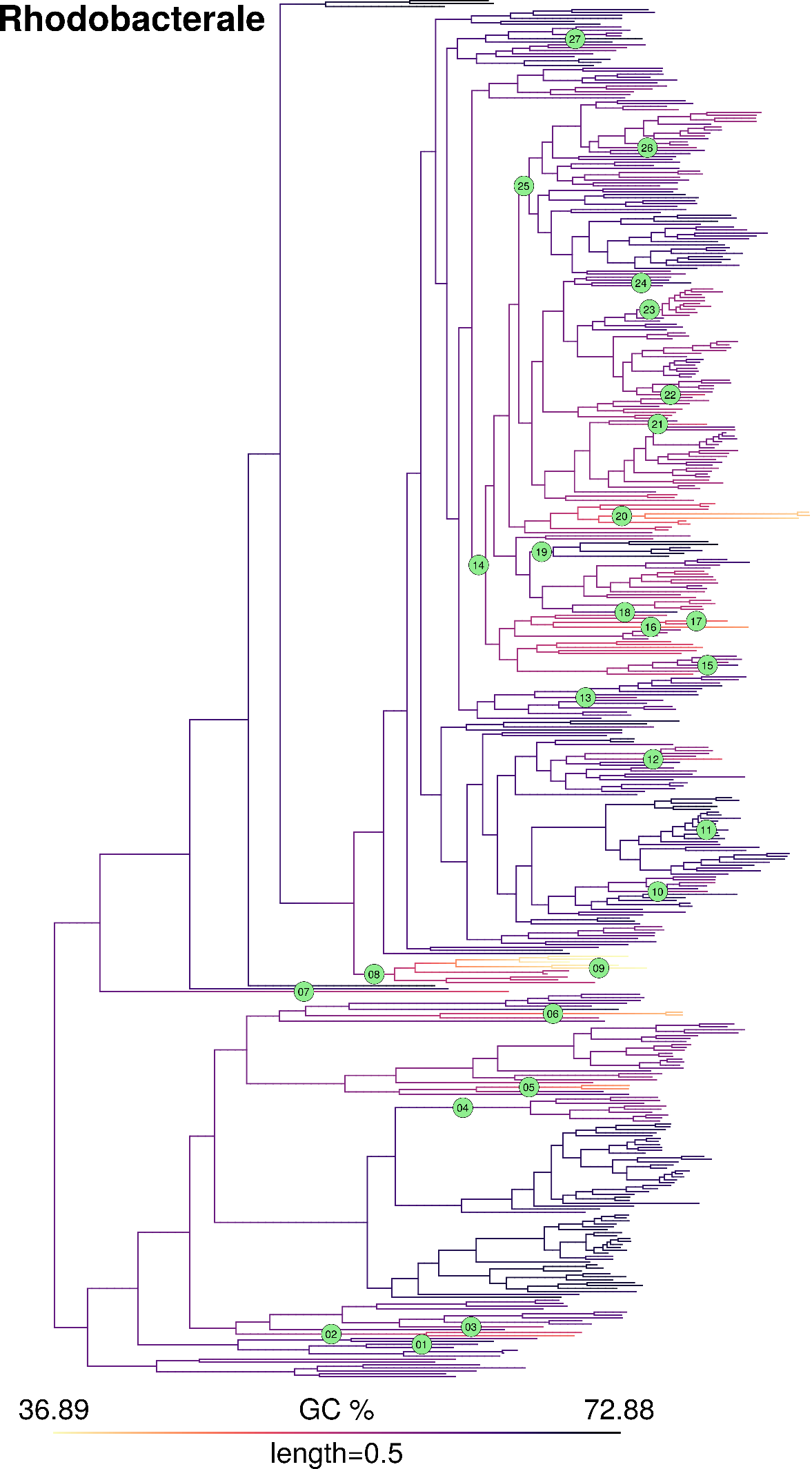

**Figure S3. GC content map and location of inferred jumps in order Rhodobacterale.**

**Figure S4. GC content map and location of inferred GC jumps in order Sphingomonadale.**

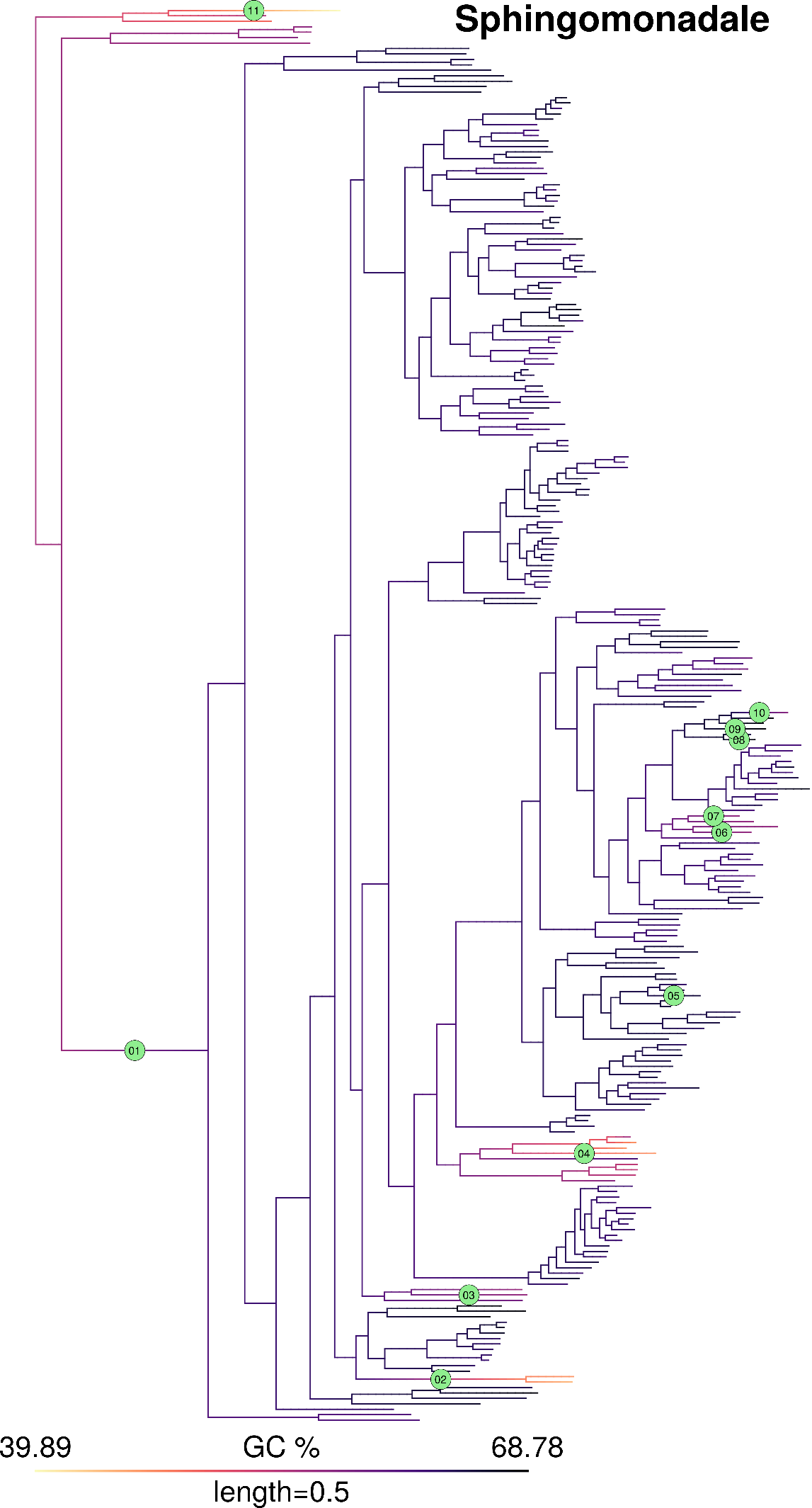

**Figure S5. GC content map and location of inferred GC jumps in order Bacteroidale.**

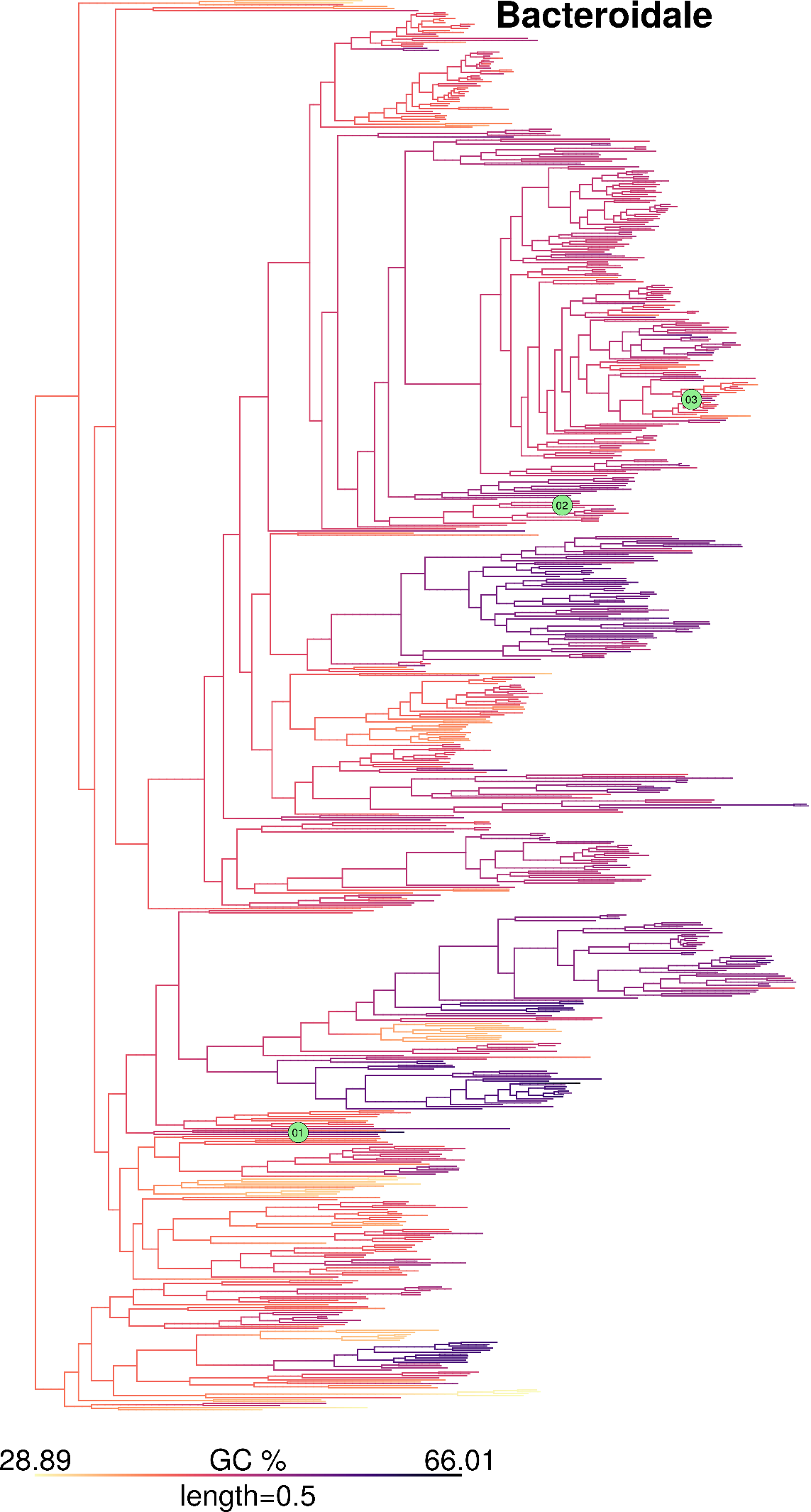

**Figure S6. GC content map and location of inferred GC jumps in order Cytophagale.**

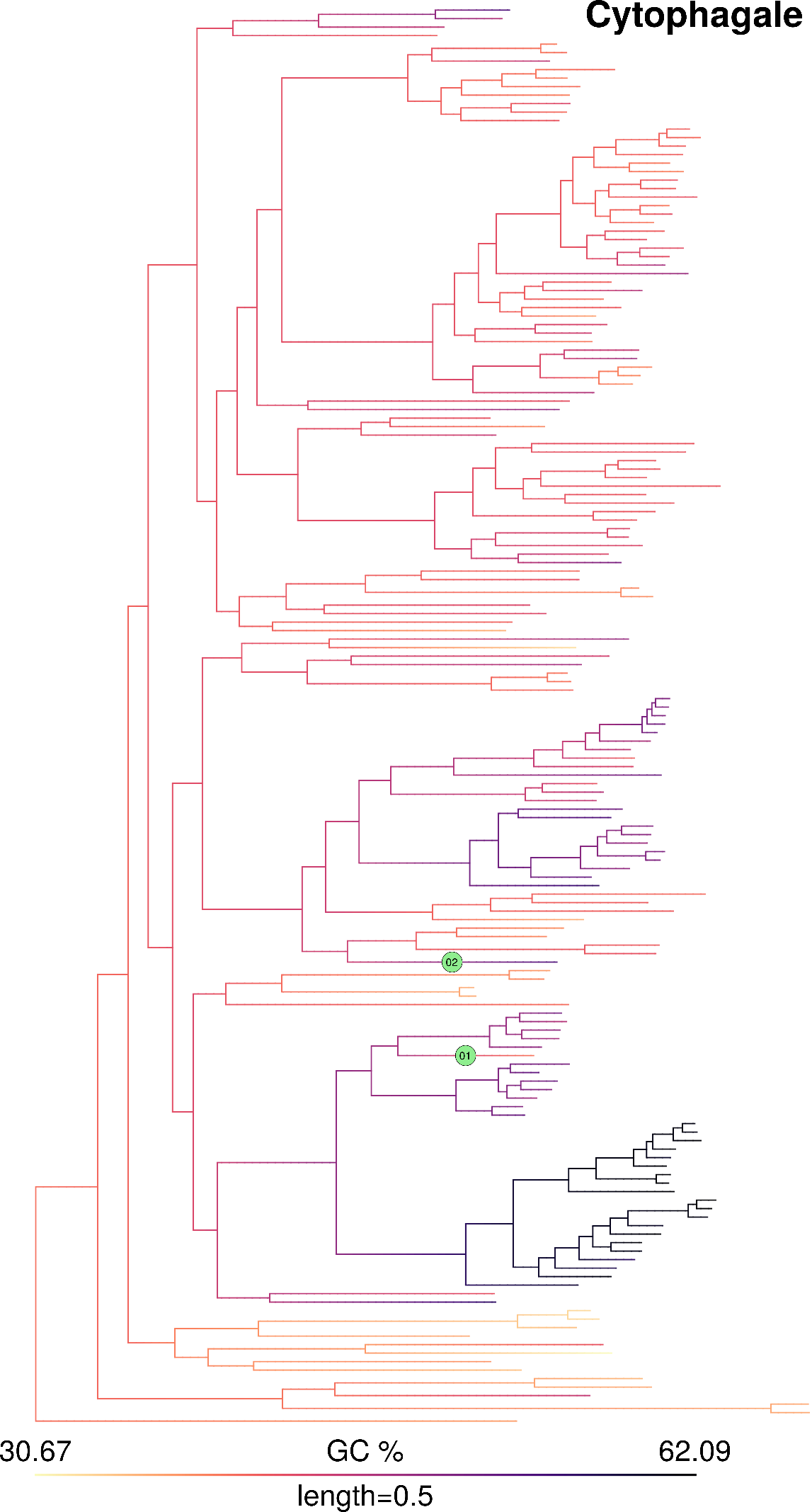

**Figure S7. GC content map and location of inferred GC jumps in order Flavobacteriale.**

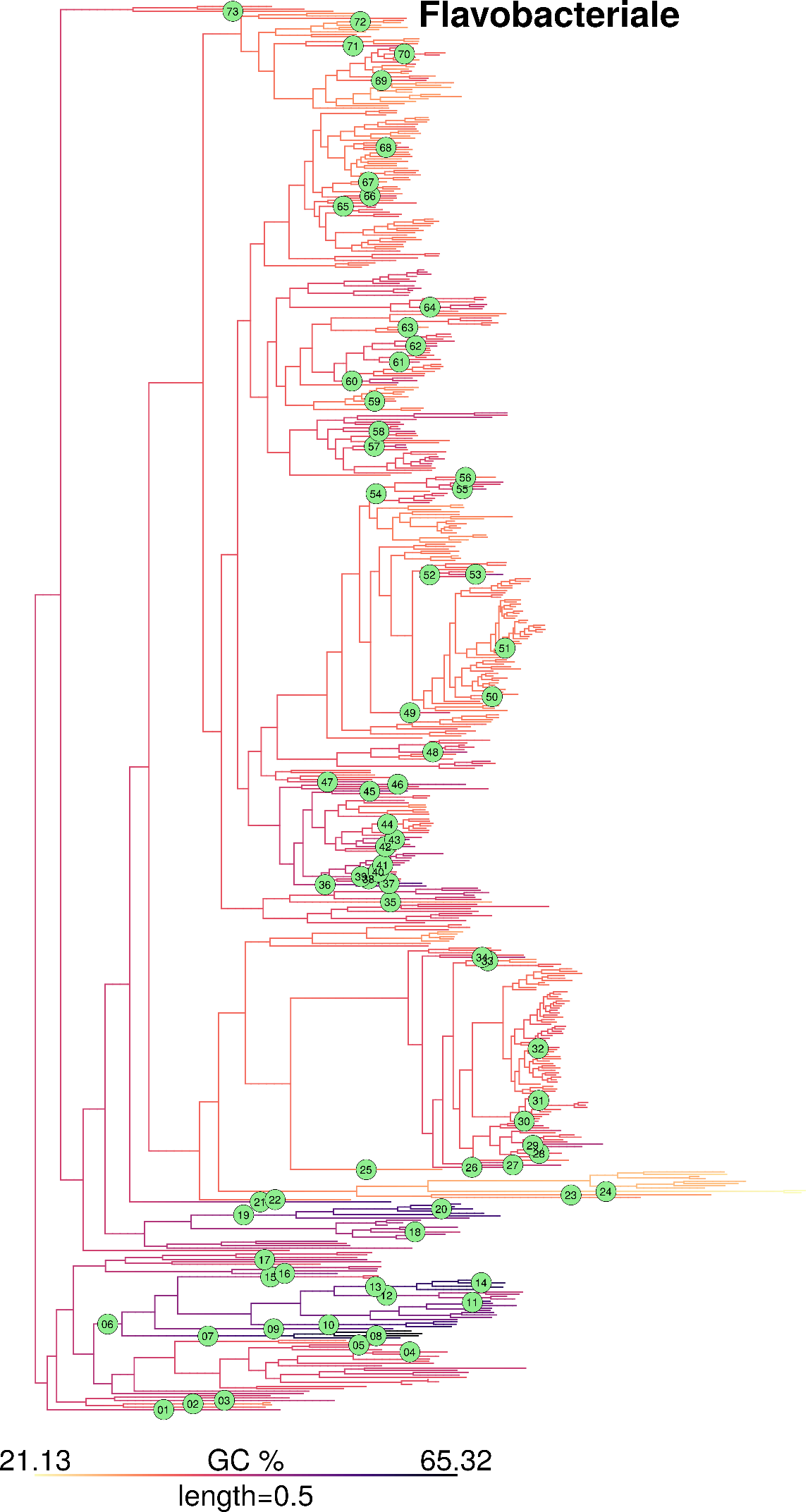

**Figure S8. GC content map and location of inferred GC jumps in order Betaproteobacteriale.**

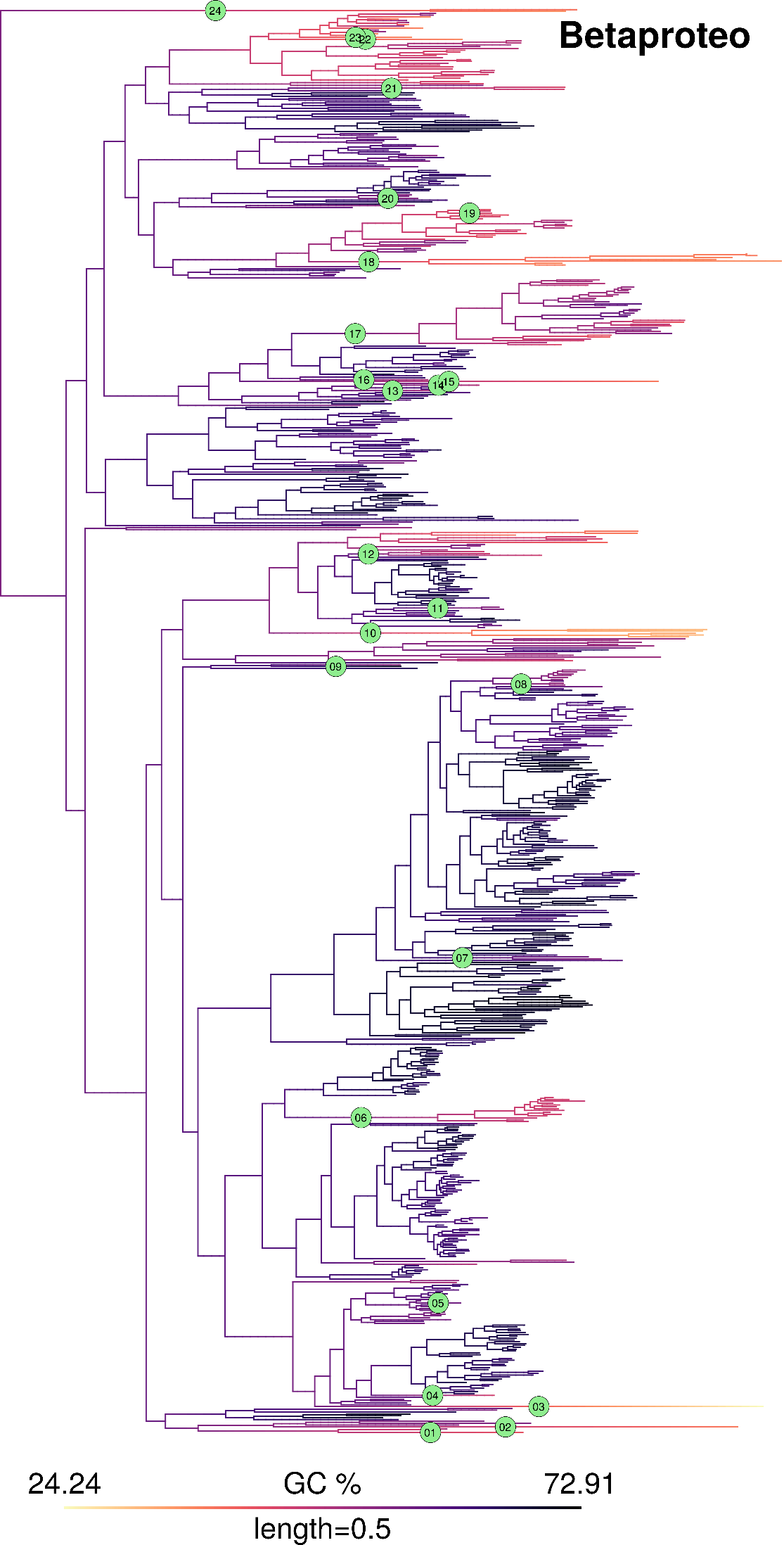

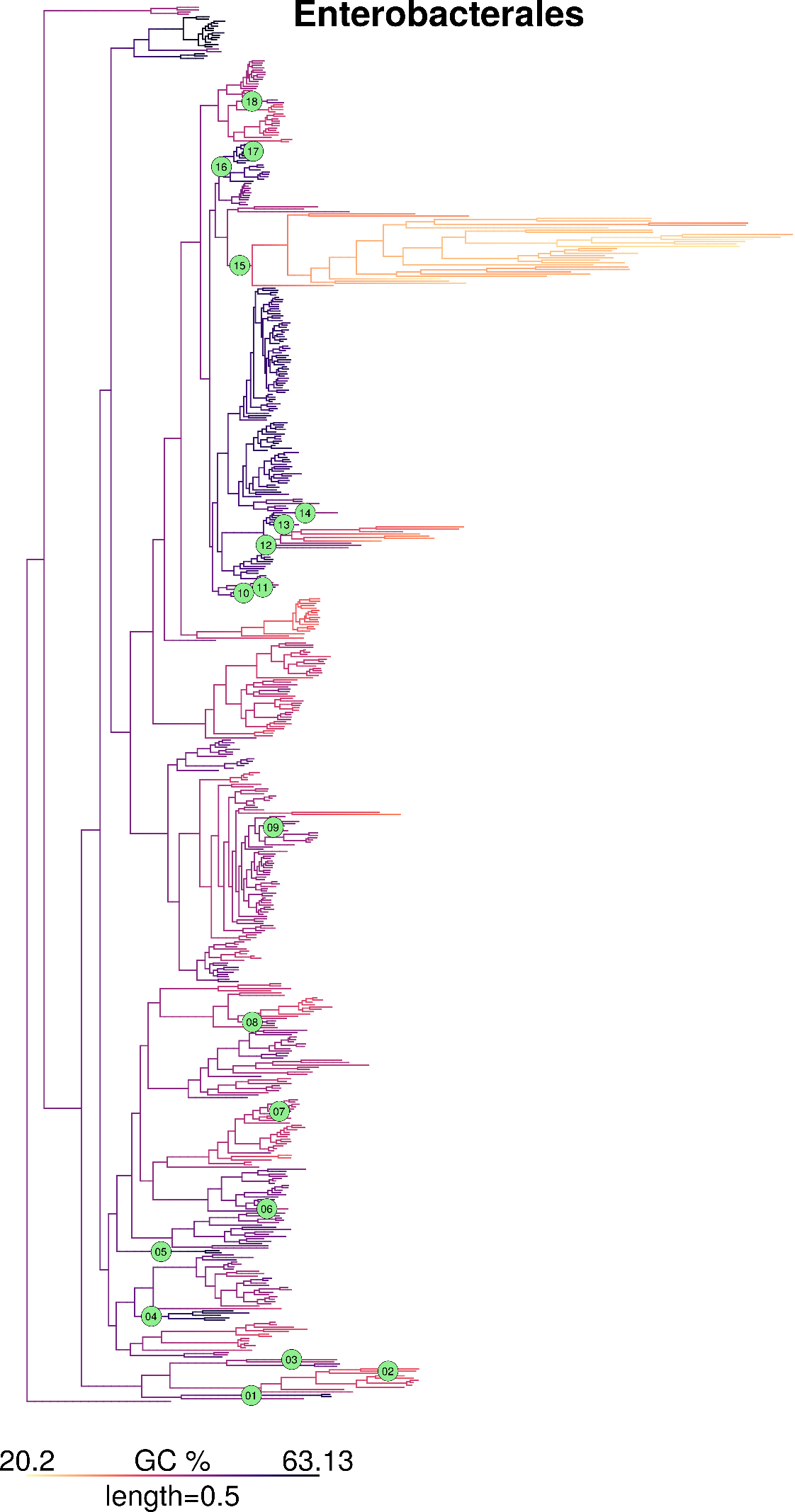
**Figure S9. GC content map and location of inferred GC jumps in order Enterobacterale.**

**Figure S10. GC content map and location of inferred GC jumps in order Pseudomonadale.**

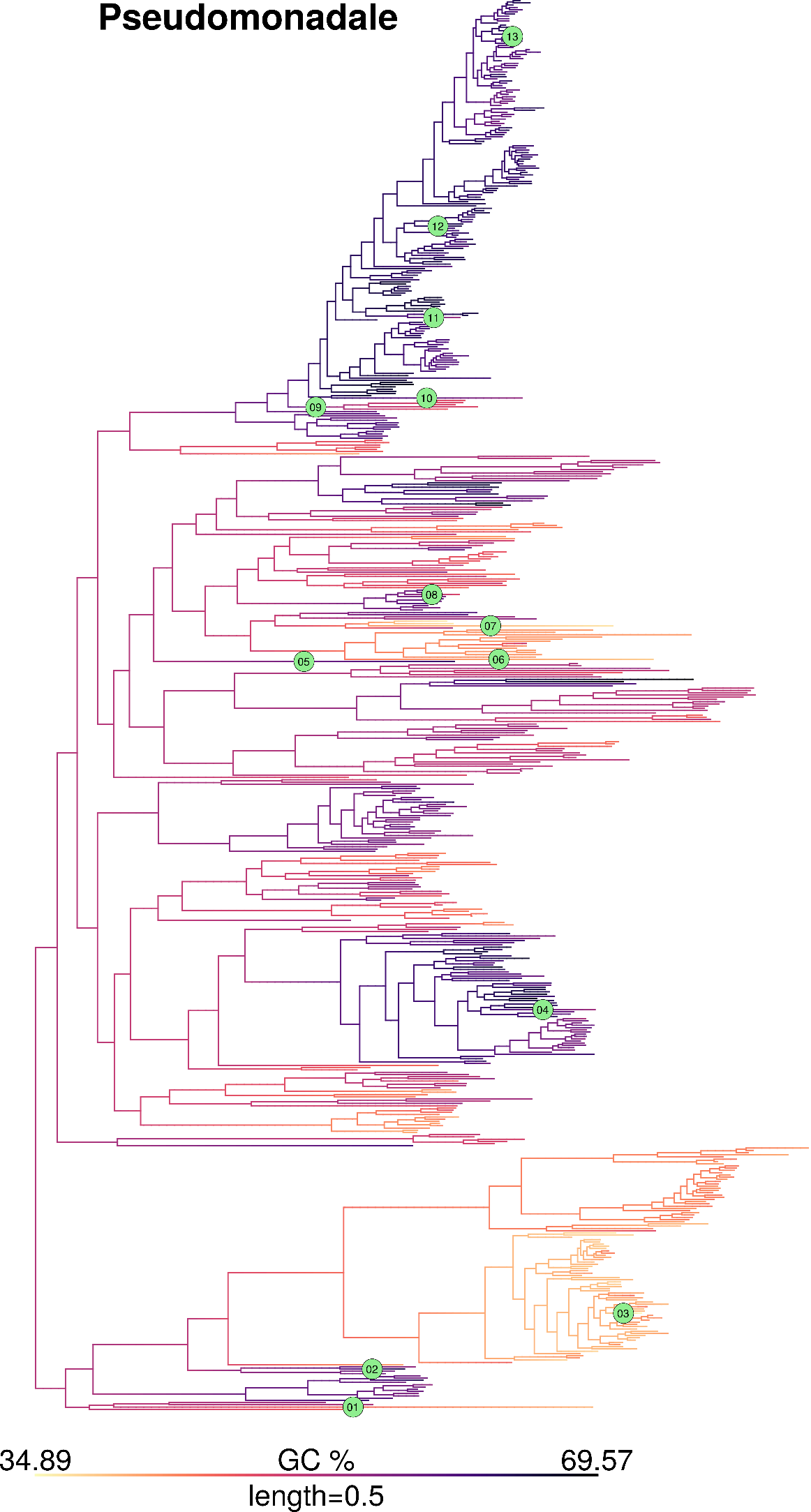

**Figure S11. Distribution of height from root of all branches, randomly placed jumps, and actually inferred jumps**

We compared the distribution of height from root of all branches, heights at which randomly placed jumps are expected, and heights of actually inferred jumps. Height for each branch is represented by the height from the root of the mid-point of the branch. Since the actual location of a GC jump within a branch is not known, we used the height of the branch on which a jump occurred as its proxy. Heights within each phylogeny were normalized by the maximum height and data was pooled across all order-level clades. (a) Distribution of normalized heights of all branches. (b) Distribution of normalized heights of randomly placed jumps. Jumps were randomly placed on branches with probabilities proportional to the branch lengths. Randomly placed jumps were simulated 10 times for each order-level clade. (c) Distribution of normalized heights of actually inferred jumps. The 1^st^ quartile, median, and 3^rd^ quartile positions (shown by dashed orange vertical lines) are not different between randomly placed and actually inferred jumps (Wilcoxon’s rank-sum test, p>>0.05).

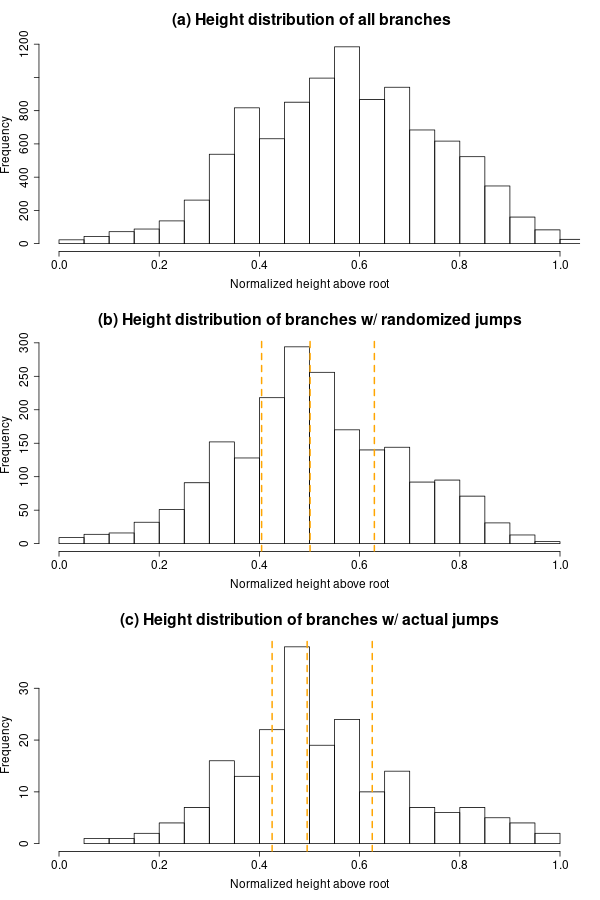

**Figure S12. Changes in GC content of ribosomal protein coding genes are strongly correlated with changes in whole genome GC content.**

We obtained the median GC content of ribosomal protein coding genes in each genome based on the annotations in “cds_from_genomic.fna” files deposited in Refseq. Further, we estimated the changes in GC content in each jump in the same way as described for whole genome GC. Data points are colored based on the magnitude of GC jump given the GC of the whole genome. The blue lines indicate regression lines with 95% confidence intervals. (A) Magnitude of jumps in GC content of ribosomal protein coding genes and genome GC content. The horizontal dashed lines shows no change in GC; whereas the sloped dashed line indicates a regression line with slope 1. The magnitude of GC jumps estimated based on ribosomal protein coding genes are lower, but correlated with those estimated from GC of the whole genome. (B) Median GC content of focal clade genomes based on ribosomal protein coding genes vs. whole genome GC content. (C) Median GC content of sister clade genomes based on ribosomal protein coding genes vs. whole genome GC content.

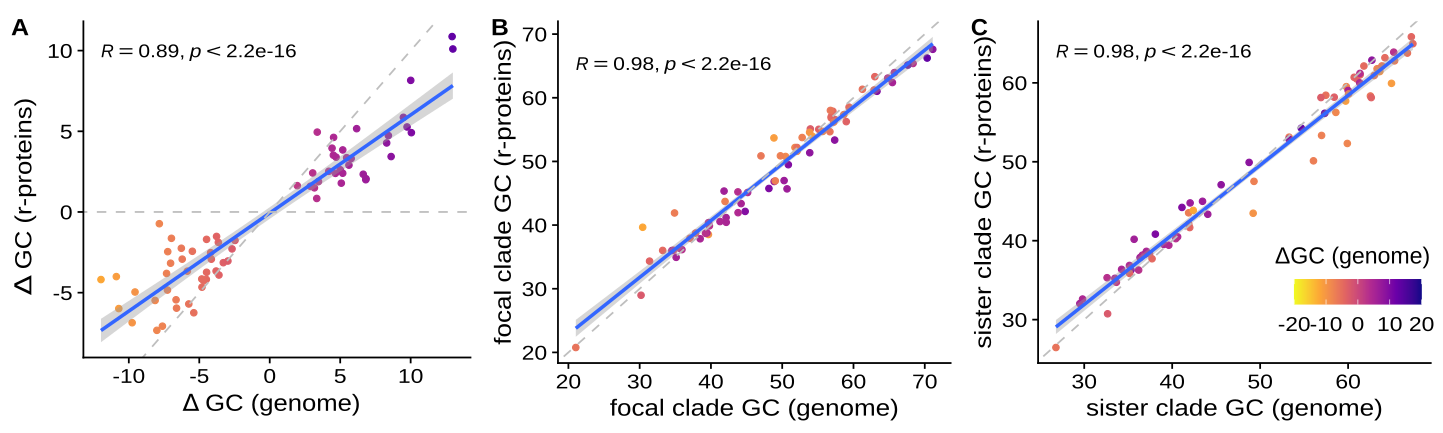
